# A new synuclein-transgenic mouse model for early Parkinson’s reveals molecular features of preclinical disease

**DOI:** 10.1101/2020.04.04.016642

**Authors:** Diana M Hendrickx, Pierre Garcia, Amer Ashrafi, Alessia Sciortino, Kristopher J Schmit, Heike Kollmus, Nathalie Nicot, Tony Kaoma, Laurent Vallar, Manuel Buttini, Enrico Glaab

## Abstract

Understanding Parkinson’s disease (PD) in particular in its earliest phases is important for diagnosis and treatment. However, human brain samples are collected post-mortem, reflecting mainly end stage disease. Because brain samples of mouse models can be collected at any stage of the disease process, they are useful to investigate PD progression. Here, we compare ventral midbrain transcriptomics profiles from *α*-synuclein transgenic mice with a progressive, early PD-like striatum neurodegeneration across different ages using pathway, gene set and network analysis methods. Our study uncovers statistically significant altered genes across ages and between genotypes with known, suspected or unknown function in PD pathogenesis and key pathways associated with disease progression. Among those are genotype-dependent alterations associated with synaptic plasticity, neurotransmission, as well as mitochondria-related genes and dysregulation of lipid metabolism. Age-dependent changes were among others observed in neuronal and synaptic activity, calcium homeostasis, and membrane receptor signaling pathways, many of which linked to G-protein coupled receptors. Most importantly, most changes occurred before neurodegeneration was detected in this model, which points to a sequence of gene expression events that may be relevant for disease initiation and progression. It is tempting to speculate that molecular changes similar to those changes observed in our model happen in midbrain dopaminergic neurons before they start to degenerate. In other words, we believe we have uncovered molecular changes that accompany the progression from preclinical to early PD.

## Introduction

The current understanding of the molecular mechanisms behind initiation and progression of Parkinson’s disease (PD) is still limited. Previous molecular studies on human post-mortem brain tissues mostly stem from late stages of PD [33, 62], making it difficult to infer causality between molecular events and pathological outcome. In addition, long post-mortem delays, several comorbidities and high inter-individual variations complicate the interpretation of molecular data. By contrast, samples from PD mouse models can be collected at any stage of the disease process, and a shared genetic background reduces inter-individual variations. Therefore, mouse models provide a useful means to investigate PD-associated molecular changes, in particular those preceding nigro-striatal degeneration. The elucidation of such early changes can shed light into disease causation and drivers of disease progression, thus in turn pointing to novel targets for intervention as well as biomarkers [16, 45].

Whole transcriptome analysis enables to compare thousands of genes between two or more groups [9], and provides a valuable method for identifying coordinated changes in gene expression patterns that are not captured by single-gene or protein measurement approaches [59]. Despite of these advantages, only a limited amount of studies have used transcriptomics on *α*-synuclein-based transgenic mouse models for PD [13, 67, 71–73, 94, 98]. Alpha-synuclein, a pre-synaptic protein believed to be a moderator of the synaptic vesicle cycle, is a major component of Lewy bodies [87]. In addition, mutations or multiplications in its genes leads to familiar forms of PD [26, 46]. Transcriptomics studies on *α*-synuclein-based transgenic mouse models for PD have only looked at differentially expressed genes (DEGs) in relation to pre-defined cellular pathways or gene sets from public annotation databases [13, 67, 71–73, 94]. Most of them used only one [71–73] or two age groups [13, 67, 94, 98] of the mouse model for their analyses, and some of them were performed in brain regions not containing dopaminergic neurons [13, 71]. Furthermore, these studies did not build custom networks from the output of the differential expression analysis by mapping each list of DEGs to a genome-scale protein-protein interaction network. An advantage of this approach is that more molecular interactions are covered than in public pathway or gene set databases. They also did not perform text mining analyses to identify putative co-occurrence associations between the human orthologues of the DEGs and Parkinson’s disease (PD) described in the literature. Finally, previous studies lack an analysis of how their findings can be translated from mouse to human biology, by systematically comparing DEGs with their human orthologues, and comparing the cellular sources of DEGs (main brain cell type where the DEGs are expressed) between mice and humans.

In this study, we characterized a new genomic *α*-synuclein transgenic model, the BAC-Tg3(SNCA*E46K) “Line 3” mice, and performed transcriptomic profiling of the ventral midbrain at 3 different ages. We found that a key factor of this model is age-dependent degeneration in the dorsal striatum, the main projection area of Substantia Nigra (SN) dopaminergic neurons [85, 104], without these neurons being lost, a phenotype reminiscent of prodromal and early PD [50]. We were interested in molecular changes driven by the *α*-synuclein transgene in the ventral midbrain at each age and across ages, and wanted to assess their disease relevance. To do this, we first identified the human orthologues of all differentially expressed genes (DEGs), then applied bioinformatical analyses consisting of (i) text mining, (ii) cellular pathway identification, and (iii) molecular sub-network analysis, both for within- and between-age comparisons. Additionally, (iv) the cellular source of the DEGs accross different brain cell types was determined for both mouse and human in order to identify similarities and differences between these two species in a PD context. Our study shows that, in this model, disease-related molecular changes at the gene expression level occur quite early, and evolve over time well before striatal degeneration reaches its peak. Our results indicate that the BAC-Tg3(SNCA*E46K) mouse line changes may provide a window into the earliest phases of PD pathology.

## Materials and Methods

### Phenotyping and tissue preparation

#### Mouse cohorts

Female mice were used at 3, 9, and 13 months of age (3M, 9M, 13M). Each age group was composed of 8 heterozygote SNCA transgene carriers (HET) and 8 wildtype (WT) littermate controls. A total of 48 female mice were used in this study.

The transgenic mouse line BAC-Tg3(SNCA*E46K), also called Tg(SNCA*E46K)3Elan*/*J or “Line 3” was originally generated by Elan Pharmaceuticals, USA, as described in detail in [30]. Briefly, the E46K mutation [103] was introduced using the Bacterial Artificial Chromosome (BAC) as described in [57]. The E46K mutation was chosen because in cultured neurons it is more toxic than the A30P and A53T mutations [39]. The mutation was confirmed by sequencing the circular BAC constructs containing the hSNCA transgene (∼ 168 Kb), which were used to perform pronuclear microinjection. Founders were identified by PCR genotyping using designated PCR primers described in [30]. Then mice were bred for over 10 generations with B6D2F1 mice (Jackson laboratories) and maintained as heterozygotes on this background, and non-TG littermates served as controls in study cohorts. For this study, mice were all females and bred in the animal facility at the Helmholtz institute Braunschweig under specific pathogen-free (SPF), climate controlled and given food and water *ad libitum*.

Genotyping by PCR (primer pair 5’-3’: forward GAT-TTC-CTT-CTT-TGT-CTC-CTA-TAG-CAC-TGG, reverse GAA-GCA-GGT-ATT-ACT-GGC-AGA-TGA-GGC) was performed on tail biopsies following the instructions found on http://www.informatics.jax.org/external/ko/deltagen/462_MolBio.html. A genotyping kit (Kapabiosystems, KR0385-v2.13) was used, following manufacturer’s instructions. Briefly, a mixture of 12.5 *µl* of “2xKAPA2G Fast Hot Start Genotyping Mix” (DNA polymerase containing mix provided with the kit mentioned in the previous section), 1 *µl* of template DNA previously prepared, 1.25 *µl* of 10 *µM* forward primer and 1.25 *µl* of 10 *µM* reverse primer was prepared in a PCR tube in a total volume reaction of 25 *µl*. The PCR products were loaded onto a 1.5% agarose gel for separation by electrophoresis, and viewed under UV after Ethidium bromide incubation.

#### Mouse brain tissue preparation and RNA extraction

All mice were euthanized in a deep anaesthesia (i.p. injection of 10 mg/kg Xylazin and Ketamine, 100mg/kg) by transcardial transfusion with PBS. After removal from the skull, brains were dissected on ice into regions, and ventral midbrains (the region enriched in dopaminergic neurons) were snap-frozen on dry ice and stored at -80°C until further processing. For RNA extraction, the RNeasy Universal Kit (Quiagen) was used, after homogenization of midbrain tissues in a Retsch MM 400 device (2 min at 22Hz, Haan, Germany). RNA concentrations and integrity were determined using a Nanodrop 2000c (Thermo Scientific) and a BioAnalyzer 2100 (Agilent), respectively. Purified RNAs were considered of sufficient quality if their RNA Integrity value was above 8.5, their 260*/*230 absorbance ratio = 1.8, and their 260*/*280 absorbance ratio = 2. RNA samples were stored at -80°C until further use.

#### Transgene expression levels by RT-PCR

##### Real Time RT-qPCR

Complementary DNA (cDNA) was prepared from 1 *µg* of total RNA, using SuperScript III reverse transcriptase (Invitrogen). A mix of RNA, oligo dT_20_ 50 *µM*, 10 mM dNTPs and sterile distilled water up to 13 *µl* was heated at 65 °C for 5 minutes, and chilled on ice for at least 1 minute prior the addition of a mixture of 5X First-Strand Buffer, 0.1 M DTT, Rnase OUT and SuperScript™ III RT (200 units/*µl*). The complete reaction was incubated at 50 °C for 60 minutes, followed by SuperScript inactivation at 70 °C for 15 minutes. Reaction tubes without Reverse Transcriptase and containing PCR water instead of RNA were included as negative controls. 80 *µl* of Rnase free water were added to the reaction, and the cDNA was placed on ice for immediate use or stored at -20 °C for future use.

Real Time RT-qPCR assays were performed using primers specific for human SNCA (Forward 5’-AAG-AGG-GTG-TTC-TCT-ATG-TAG-GC-3’; Reverse 5’-GCT-CCT-CCA-ACA-TTT-GTC-ACT-T-3’), murine Snca (Forward 5’-CGC-ACC-TCC-AAC-CAA-CCC-G-3’; Reverse 5’-TGA-TTT-GTC-AGC-GCC-TCT-CCC-3’), or total synuclein (Forward 5’-GAT-CCT-GGC-AGT-GAG-GCT-TA-3’; Reverse 5’-GCT-TCA-GGC-TCA-TAG-TCT-TGG-3’), with GAPDH (Forward 5’-TGC-GAC-TTC-AAC-AGC-AAC-TC-3’; Reverse 5’-CTT-GCT-CAG-TGT-CCT-TGC-TG-3’) as an internal house-keeping transcript. The qPCR reaction contained 2 *µl* of cDNA, 10 *µM* forward and reverse primers, 1X iQ*™* SYBR®Green Supermix (Bio-Rad) and PCR grade water up to a volume of 20 *µl*. Each qPCR reaction was run in triplicates on a LightCycler®480 II (Roche). The thermo cycling profile included an initial denaturation of 3 minutes at 95°C, followed by 40 Cycles of 95°C for 30 seconds, 62°C for 30 seconds and 72°C for 30 seconds with fluorescent data collection during the 62°C step. Data acquisition was performed by LightCycler®480 Software release 1.5.0 SP4, version 1.5.0.39. Expression values for each target gene were normalized to the housekeeping gene GAPDH (Δ*Ct*), and the results are shown as fold change in expression (2^−Δ*Ct*^).

#### Immunostaining procedure

Immunofluorescence was performed using standard procedures. Fifty micron free-floating sections were washed (2 × 5 min) under circular agitation in washing buffer (PBS with 0.1% TritonX100), then permeabilized for 30 min in PBS containing 1.5% TritonX100 and 3% hydrogen peroxide (Sigma; #31642). Then, sections were washed again in washing buffer (2 × 5 min) before being incubated in the same buffer with 5% bovine serum albumin (BSA, Sigma, A7030) as blocking solution for 30 min. After being rinsed (2 × 5 min), sections were incubated with primary antibodies (mouse anti-human synuclein, Sigma S5566 (clone SYN211) for human alpha-synuclein; rabbit anti-synuclein, Sigma S3062, for both human and mouse alpha-synuclein) diluted 1:1000 in washing buffer supplemented with 0.3% Triton X-100, 2% BSA, overnight with shaking at room temperature. After 3 more washing steps (10 min each), sections were incubated with appropriate secondary antibodies (goat anti-rabbit coupled to Alexa-488, Invitrogen A11034; or goat anti mouse coupled to Alexa-488, Invitrogen A21121), diluted 1:1000 in washing buffer, in the dark for 2 hours at room temperature. Next, the sections were washed (3 × 10 min) in washing buffer, and once in TBS (Trizma base 6.06g.L-1, 8.77 g.L-1 NaCl, adjusted with HCl to pH7.4). Finally, sections were mounted on glass slides, coverslipped using Dako fluorescent mounting medium (DAKO NA; #S3023), and imaged, at least 24h later, using a Zeiss LabA1 microscope, coupled to a Zeiss Axiocam MRm3 digital camera, and to a PC using the Zeiss Zen Blue 2012 software.

#### Quantitation of nigro-striatal degeneration

Immunofluorescence staining for tyrosine hydroxylase (TH) and dopamine transporter (DAT) was performed using standard procedures as described above. Sections were double-stained for both markers. Primary antibodies (chicken anti-TH, Abcam #76442; Rat anti DAT, Millipore MAB369) were diluted 1:1000 in washing buffer containing 0.3% Triton X-100, 2% BSA, and incubated with sections overnight, with shaking, at room temperature. Secondary antibodies (goat anti-chicken coupled to Alexa 568; goat anti-rat coupled to Alexa 594 Invitrogen A11007), diluted 1:1000 in washing buffer, were incubated with sections in the dark for 2 hours at room temperature. Then, sections were mounted on Superfrost glass slides, coverslipped using Dako fluorescent mounting medium (DAKO; #S3023), and imaged at least 24h later. Imaging was done using a Zeiss AxioImager Z1 upright microscope equipped with a PRIOR motorized slide stage, coupled to a “Colibri” LED system to generate fluorescence light of defined wavelengths, and an Mrm3 digital camera for image capture. Striatal pictures were acquired using the apotome module at 40X magnification (pictures size: 223×167 *µ*m), and SN were acquired at 10X magnification (picture size: 895×670 *µ*m) with regular epifluorescence. The Zeiss imaging system was controlled by Zeiss’ Blue Vision software. To determine neurodegeneration in the striatum [64], 2-3 striatal sections per animal were imaged, and 3 pictures from the dorsal striatum were collected for each section. Images were converted to Tiff image. Then, after thresholding, the percent image area occupied by TH or DAT-positive structures were measured using the publicly available imaging software FIJI [81]. Measuring of relative levels of human *α*-syn and its correlation with loss of TH-positive axons in the striatum was done on TH/*α*-syn double-labeled sections. For all images used for quantitation of TH, a parallel quantitation of *α*-syn was performed. Since the images were 8-bit images, the relative level of *α*-syn was measured, using the same imaging software, by determining the mean grey value, on a scale of 0-255, of *α*-syn positive synapses, similar to the approach described in [12]. Degeneration of SNs was assessed as described in the supplementary material in Ashrafi et al. [4]. Briefly, 895×670 *µ*m pictures of the whole SNs were sampled each 4th section throughout the whole ventral midbrain, and each picture was assigned to one of the four SN typical subregions. The selected four pictures from the four subregions from each mouse were summed together to calculate the “Cumulated surface” used for the quantification. This method has been validated by stereological assessment of the total number of TH-positive neurons in the SN. All individual values for the different measurements obtained for each mouse were averaged to give one value for each mouse. All quantitation measurements were performed on blinded sections and codes were not broken until the measurements were completed.

Statistical analyses were done using the GraphPad Prism version 8.0.0 for Windows, GraphPad Software, San Diego, California USA. Normality of data distribution was tested with Kolmogorov-Smirnov normality test. All datasets analyzed showed a normal distribution, thus two-way ANOVA (for two independent variables, in this case genotype and age), followed by Tukey’s or Sidak’s test post-hoc, was used. Significance of correlation was tested by Pearson’s test. A p-value below 0.05 was considered significant for all analyses.

#### Microarray analysis

RNA purity and integrity were re-confirmed NanoDrop1ND-1000 spectrophotometer and Agilent 2100 Bioanalyzer with RNA 6000 Nano assay kit. Only RNAs with no sign of marked degradation (RIN > 7-8) were considered good quality and used for further analysis. GeneChip Mouse Gene 2.0ST Arrays (Affymetrix) were used. Total RNAs (150 ng) were processed using the Affymetrix GeneChip^®^ WT PLUS Reagent Kit according to the manufacturer’s instructions (Manual Target Preparation for GeneChip^®^ Whole Transcript (WT) Expression Arrays P/N 703174 Rev. 2). In this procedure, adapted from [11], the purified, sense-strand cDNA is fragmented by uracil-DNA glycosylase (UDG) and apurinic/apyrimidinic endonuclease 1 (APE 1) at the unnatural dUTP residues and breaks the DNA strand. The fragmented cDNA was labelled by terminal deoxynucleotidyl transferase (TdT) using the Affymetrix proprietary DNA Labelling Reagent that is covalently linked to biotin; 5.5 *µ*g of single-stranded cDNA are required for fragmentation and labelling, then 3.5 *µ*g of labeled DNA and hybridization controls were injected into an Affymetrix cartridge. Microarrays were then incubated in the Affymetrix Oven with rotation at 60 rpm for 16 hr at 45°C, then the arrays were washed and scanned with the Affymetrix^®^ GeneChip^®^ Scanner 3000, based on the following protocol: UserGuide GeneChip^®^ Expression Wash, Stain and Scan for Cartridge Arrays P/N 702731 Rev. 4, which generated the Affymetrix raw data CEL files containing hybridization raw signal intensities. All microarray analyses were conducted in the same run as one batch.

### Pre-processing and quality control

Probe correction, removal of array-specific background noise and normalization were performed using the SCAN pre-processing procedure [74], implemented in the Bioconductor [31, 38] SCAN.UPC package (version 2.24.1).

The Bioconductor arrayQualityMetrics package (version 3.38.0) was used for quality control. Samples were removed if at least two of the three following quality control tests failed: outlier test of distances between arrays, boxplots, MA plots (i.e. the scatter plots of the distribution of the log2 expression ratio (M) against the average expression signal (A)) [47].

### Bioinformatics tools and statistical analyses

#### An overview

The workflow for our analyses is depicted in Figure 1. After pre-processing and quality control, differentially expressed genes (DEGs) between genotypes for each age group (HET versus WT; genotype-dependent DEGs), as well as DEGs between age groups for each genotype separately (3M versus 9M, 9M versus 13M; age-dependent DEGs) were determined. We used the resulting lists of DEGs for text mining, molecular pathway, gene set and network analysis. We also determined the cellular origin of the DEGs and of their human orthologues using public databases.

**Figure 1.**
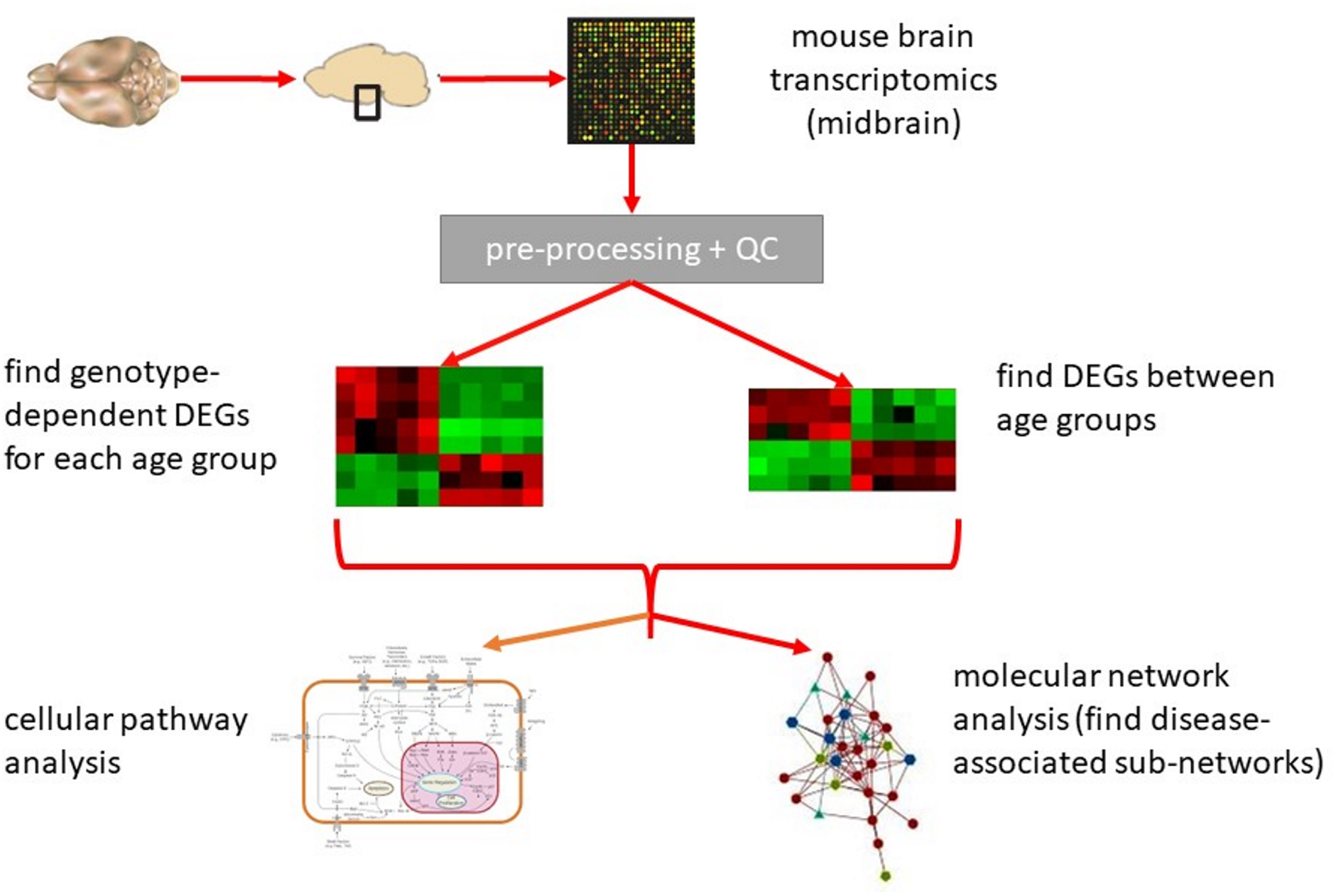
Data analysis workflow.

The term *“differentially expressed gene”*, or DEG, is used to describe differential mRNA expression in microarrays. It’s important to keep in mind that these measurements not only reflect gene transcriptional activity, but also factors as mRNA stability and degradation. *Pathways* describe biological phenomena, such as gene regulation, signal transduction and metabolic processes, in terms of their interactions between biological entities (genes, proteins, metabolites). They are represented by a graph consisting of nodes (biological entities) and edges (interactions). In contrast, a *gene set* is a collection of genes involved in the same biological phenomenon, and does not use any information about interactions. This study will focus on gene sets representing *biological processes from the Gene Ontology (GO) collection.*

*Molecular network analysis* is complementary to pathway analysis as these methods do not rely on pre-defined pathways or gene sets. These tools build a new *molecular sub-network* based on known molecular interactions from a genome-scale protein-protein interaction network extracted from public databases. As molecular interaction databases cover more interactions than pathway- or gene set databases, molecular network analysis has the potential to provide novel insights on complex interactions (represented graphically by edges) between biological entities (represented by nodes).

#### Differentially expressed genes (DEGs)

To perform differential expression analysis on the microarray data, pairwise comparisons between groups were conducted with the empirical Bayes moderated t-statistic [84] implemented in the Bioconductor limma package (version 3.38.3) [77], with adjustment for multiple testing using the Benjamini-Hochberg procedure [5](false discovery rate (FDR) ≤ 0.05). Since small activity changes in regulatory genes may have important downstream effects [55], we included all genes with statistically significant differential expression (FDR ≤ 0.05), independent of the effect size of the change i.e. without applying a log fold-change based threshold.

- To identify *genotype-dependent DEGs*, we performed pairwise comparisons between the two genotypes (WT and HET) at each age (3M, 9M, and 13M).
- To identify *age-dependent DEGs* that are specifically associated with synuclein transgene expression in HET mice, we first performed pairwise comparisons between consecutive age-groups (9M versus 3M, 13M versus 9M) for each genotype separately (WT and HET). Since 9M was the middle age group, we picked this age group to compare to the younger (3M) and older (13M) group, for both groups separately. We selected the genes that were differentially expressed for both age-group comparisons. Then, we removed age-related DEGs found in both WT and HET from the totality of those found in HET mice, and used the remaining DEGs for pathway, gene set and molecular network analysis. With this approach, we observed three patterns of gene expression: genes whose expression gradually increased from 3M to 9M and then on from 9M to 13M (type 1), those whose expression gradually decreased from 3M to 9M and then on from 9M to 13M (type 2), and those switching in expression directionality between 3M to 9M and 9M to 13M (type 3). This last type of DEGs could be subdivided in two expression patterns: genes whose expression increased between 3M and 9M, then decreased between 9M and 13M (Type 3a), and genes whose expression decreased between 3M and 9M, then increased between 9M and 13M (Type 3b).

Significance of the overlap of the DEGs between two comparisons was determined using Fisher’s exact test.

The DEGs identified were further paired down with the help of successive translational relevance assessment steps.

#### Human orthologues of DEGs

In a first relevance assessment step, human orthologues for all DEGs were identified and extracted from Ensembl Biomart database (https://www.ensembl.org/biomart). Orthologues in Ensembl Biomart are determined from gene trees, constructed using one representative protein per gene for each species. The tree is based on sequence alignment, and speciation nodes in the tree correspond to orthologues. These orthologues datasets were the ones used for text mining, pathway, gene-set, network and cellular abundance analyses.

Some of the DEGs with human orthologues in Ensembl BioMart, were identified as mouse specific in the database NCBI Homologene (https://www.ncbi.nlm.nih.gov/homologene). Table S1 presents an overview of these genes, and also shows expression levels in the brain according to the Human Proteome Atlas (https://www.proteinatlas.org/ENSG00000069493-CLEC2D/tissue).

#### Cellular source for the DEGs

In a second relevance assessment step, for all DEGs, the brain cell type where the genes show the highest expression was extracted from the Brain RNAseq database (https://www.brainrnaseq.org/) [106] for both the mouse DEGs and their human orthologues. In case of expression in different cell types, the cell type having the highest Transcripts Per Million score (TPM) for the studied gene was considered as the cell type of expression.

#### Text mining and PMI score

In a third relevance assessment step, to identify and rank putative functional associations between DEGs and Parkinson’s disease (PD) previously described in the literature, a text mining analysis using the Pointwise Mutual Information (PMI) co-occurrence measure was performed [91]. The relative co-occurrences of the human orthologues of the DEGs for the comparisons listed above with the controlled vocabulary Medical Subject Headings (MeSH) term “Parkinson disease” were quantified in full-text papers from PubMed. For each gene in the list of DEGs, the PMI score is calculated as follows:

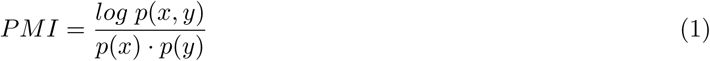

where p(x) is the proportion of full-text papers in PubMed that match the gene, p(y) is the proportion of full-text papers in PubMed that match the MeSH term “Parkinson Disease”, p(x,y) is the proportion of full-text papers in PubMed that match the gene and the MeSH term “Parkinson Disease”. PMI calculates how much more the gene and the MeSH term “Parkinson Disease” co-occur than expected by chance. Genes with a higher PMI score display more frequent relative co-occurrences with the MeSH term “Parkinson disease” in relation to the occurrences of the individual terms in PubMed, pointing to candidate gene-disease associations for which specific literature references can be retrieved (see Results section).

#### Comparability of the mouse model to human data

In a fourth relevance assessment step, to test the relevance of the mouse model to study molecular events leading to an early-PD like phenotype, the human orthologues of the lists of DEGs between genotypes (HET versus WT) obtained from the mouse data for each age (3M, 9M, 13M) were compared to the list of DEGs from a recent meta-analysis of transcriptomics data from post-mortem brain samples of PD patients and controls conducted by Kelly et al. [48]. Thus, with these 4 relevance assessment steps, we evaluated whether the mouse model data can provide an adequate reflection of the alterations associated with early PD progression.

#### Pathway and gene set analysis

To detect molecular pathway and gene set alterations associated with the progression of PD-like pathology in HET mice, an enrichment analysis was conducted using the web-based software GeneGO MetaCore*™* (https://portal.genego.com/). The input table was the list of human orthologues of the genes identified as DEGs in the mouse dataset, with adjusted p-value ≤ 0.05, together with the logarithmic fold change to determine the expression directionality. Because small-effect size changes in transcription factors may be relevant for the downstream analyses [55], we did not apply a log fold change threshold. In case several mouse genes had the same human orthologue, the gene having the highest average expression was taken as representative. The p-values for the pathways were corrected for multiple hypothesis testing using the approach by Benjamini and Hochberg [5] and an FDR threshold of 0.05 was applied. Only pathways with size between 5 and 500 network objects (gene products), including at least 3 network objects from the data (DEG list), were considered. Furthermore, pathways whose function is not known in the brain, but only in other organs, were removed.

To keep in line with the analysis of DEGs that had different patterns of expression changes (see above), a separate pathway and gene set analysis was performed for:

- Genotype-dependent changes: the DEGs between HET and WT, within each age group.
- Age-dependent changes in HET only: genes whose expression increases with age (type 1), genes whose expression decreases with age (type 2); genes whose expression switches directionality (type 3a: increased between the 3M and 9M, decreased between 9M and 13M; type3b: decreased between the 3M and 9M, increased between 9M and 13M). Age-dependent pathways and gene sets that were identified by this approach were named the same way as the DEGs they were based on (type 1, type 2, type 3a and b), based on whether they were upregulated (type 1), downregulated (type 2), or switching their regulation directionality as the HET mice aged (type 3a and 3b)

#### Network analysis

Network analysis investigates altered interactions between key components of different pre-defined pathways or biological processes. It does so by mapping each list of DEGs to a genome-scale protein-protein interaction network. Analysis of the custom networks build in this way, called molecular sub-networks, provides complementary information to the pathway analysis on associations between gene sets and biological processes.

In our study, molecular sub-networks were determined by analyzing the lists of DEGs that were used for the pathway analyses with the GeneGO MetaCore*™* default “Analyze network”(AN) procedure. Using information on molecular interactions and canonical pathways, the procedure identifies and ranks altered molecular sub-networks surrounding the mapped gene products corresponding to the DEGs. The resulting molecular sub-networks include a number of additional proteins and protein complexes that interact directly or indirectly with the gene products of DEGs. This algorithm first builds a large network by gradually expanding a network around every gene product from the DEGs, then it gives priority to proteins and protein complexes having the highest connectivity (incoming and outgoing edges) with the initial protein or protein complex. The larger network is divided into a maximum of 30 sub-networks of maximum size 50 (default settings), so that each edge only occurs in one sub-network. Nodes may be part of different sub-networks but connected by different edges. A p-value for the intersection of the gene products from the list of DEGs and the nodes in the sub-network was calculated by Fisher’s exact test. Only networks including at least 3 gene products of the input list were selected.

## Results

### Transgene expression levels in BAC-Tg3(SNCA*E46K) “Line 3” mice

The levels of SNCA transgene RNA, of murine Snca RNA, and of total RNA from both (murine and human), all measured by RT-qPCR, for the ventral midbrain, are shown in Figure 2A. Transgene SNCA RNA was detected in all mice that tested positive for the transgene by genotyping (HET mice), but not in wildtype (WT) littermates. Levels of murine Snca RNA were not significantly different between mice of the two different genotypes. Levels of total alpha-synuclein RNA indicate an approximately 1.8-fold transgene expression in HET mice compared to WT littermates. Taken together, this indicates that the increased overall *α*-syn RNA expression in HET was due to transgene. Alpha-synuclein protein, human or total protein, were revealed by immunofluorescence (Figure 2B). Transgenic human alpha-synuclein was localized mainly in the neocortex, hippocampus, thalamus, striatum and midbrain areas (including SN), and its distribution roughly matched that of endogenous alpha-synuclein in WT mice.

**Figure 2.**
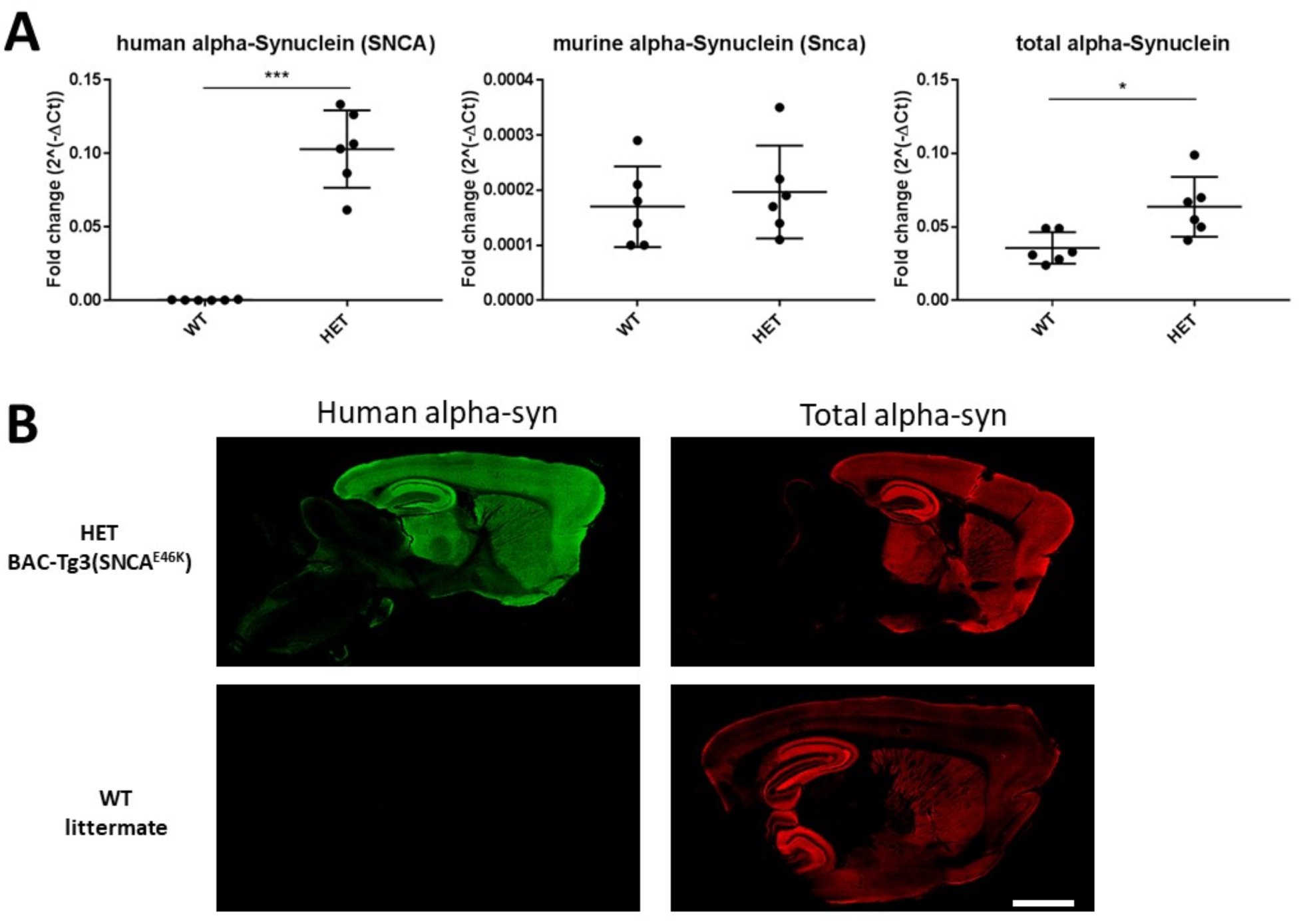
Alpha-synuclein expression in BAC-TG3(*SNCA*^*E*46*K*^) “Line 3” mouse brains. RT-PCR measurements (A) showed: that SNCA was detected in all HET mice, that the levels of RNA for murine Snca were similar between HET mice and WT littermates, and that the levels of overexpression (measured by levels of total RNA for both SNCA and Snca) were about 1.8-fold (78%) in ventral midbrain of HET mice compared to that of WT littermates. \*\*\**p* < 0.0001, \**p* < 0.05 by unpaired Student’s T test. Fluorescent immunostaining showed that the transgene human alpha-synuclein (panels in left column) shows roughly the same regional distribution than the endogenous murine alpha-synuclein (panels in right column). Scale bar (B) = 1.4 mm.

### Age-dependent neurodegeneration in the dorsal striatum but not in the Substantia Nigra

To determine *α*-syn transgene induced effects in the nigro-striatal circuit in HET mice at different ages, we measured the integrity of TH-positive axons and DAT-positive synapses in the dorsal striatum, and estimated the number of TH-positive neurons in the SN in HET and WT control mice.

Our results are shown in Figure 3. We found an age-dependent decline of TH-positive axons (Figure 3A) and in DAT positive synaptic terminals (Figure 3B) in HET mice. We did not observe loss of TH-positive neurons in the SN (Figure 3C) in these mice. Importantly, striatal loss of TH-positive axons in 13M old HET mice significantly correlated negatively with relative levels of transgenic *α*-syn (r = -0.94, p = 0.017, by Pearson’s). This indicates that degeneration of TH-positive axons in the dorsal striatum is caused by human *α*-syn, and that the neurodegenerative phenotype of BAC-Tg3(SNCA^*E*46*K*^) mice is not a consequence of a transgene insertion artefact. Because striatal degeneration without measurable loss of TH-positive nigral neurons is a feature of incipient Lewy Body disease [18, 21] and early PD [50], we decided to use our model as a tool to study gene expression changes in the ventral midbrain leading up to the degeneration in the dorsal striatum.

**Figure 3.**
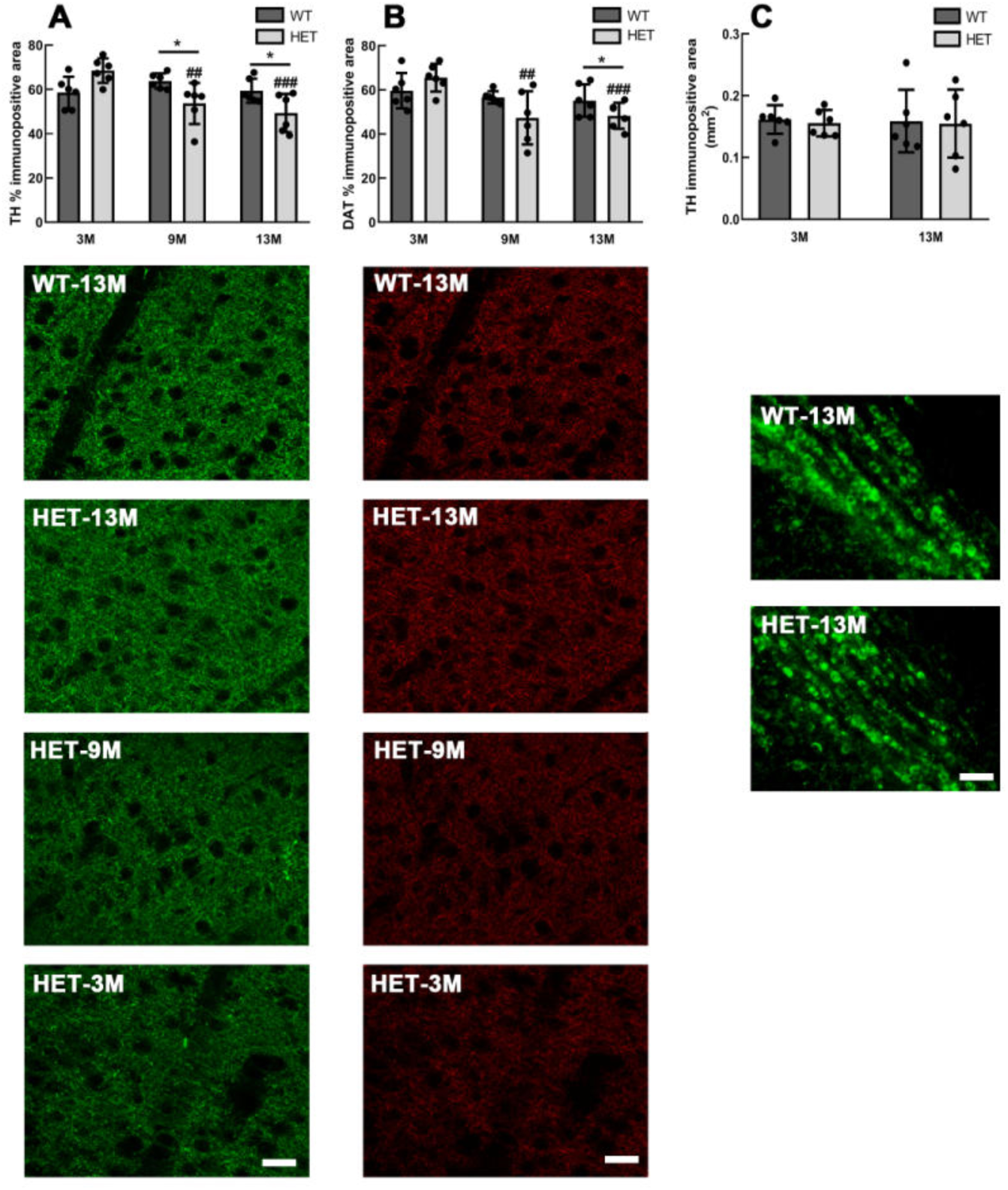
Age-dependent degeneration in the dorsal striatum but not the Substantia Nigra, in heterozygous (HET) BAC-Tg3(SNCA^*E*46*K*^) mice compared to littermate wildtype (WT) controls. Measurements for Tyrosine-hydorxylase (TH) and Dopamine Transporter (DAT) were performed as described in Material and Methods. Two-way ANOVA followed by post-hoc tests were used for statistical analysis. N= 6 mice per age-group and genotype. ^∗^ symbol for p<0.05 in genotype comparisons, ## symbol for p<0.01 and ### for p<0.001 when comparing 9M HET and 13M HET mice, respectively, to 3M HET mice. Scale bars: 30 *µ*m for immunostaining panels in columns A and B, and 60 *µ*m for immunostaining panels in column C.

### Uncovering and assessing the relevance of genotype-dependent and age-dependent DEGs

Table 1 shows the number of differentially expressed genes (DEGs) for each comparison used in this study to uncover genotype- and age-dependent DEGs, together with the corresponding number of human orthologues according to the public repository Ensembl BioMart (https://www.ensembl.org/biomart).

**Table 1.** Number of differentially expressed genes (DEGs) identified for the different comparisons, together with the corresponding number of human orthologues according to Ensembl BioMart ((https://www.ensembl.org/biomart). ^∗^: 2 genes are identified as ‘only expressed in mice’ in NCBI Homologene (https://www.ncbi.nlm.nih.gov/homologene)(see Table S1); ^∗∗^: 3 genes are identified as ‘only expressed in mice’ in NCBI Homologene (see Table S1); ^∗∗∗^: 10 genes are identified as ‘only expressed in mice’ in NCBI Homologene, 4 of them have no reported brain expression, based on Human Protein Atlas (https://www.proteinatlas.org/ENSG00000069493-CLEC2D/tissue) (see Table S1).

| comparison | number of mouse DEGs | number of human orthologues |
| --- | --- | --- |
| 3M HET vs 3M WT | 741 | 584 |
| 9M HET vs 9M WT | 7 | 5 |
| 13M HET vs 13M WT | — | — |
| 9M HET vs 3M HET | 1986 | 1401* |
| 13M HET vs 9M HET | 1837 | 1532** |
| 9M WT vs 3M WT | 6574 | 4526*** |
| 13M WT vs 9M WT | 2002 | 1664* |

We first identified genotype-dependent (HET mice versus WT mice) DEGs at each age. The approach is detailed in the Materials and Methods section. We observed 584 DEGs in 3M mice, 5 in 9M mice, and none in 13M mice. There was no overlap between the DEGs in 3M mice and those in 9M mice (Figure 4A).

**Figure 4.**
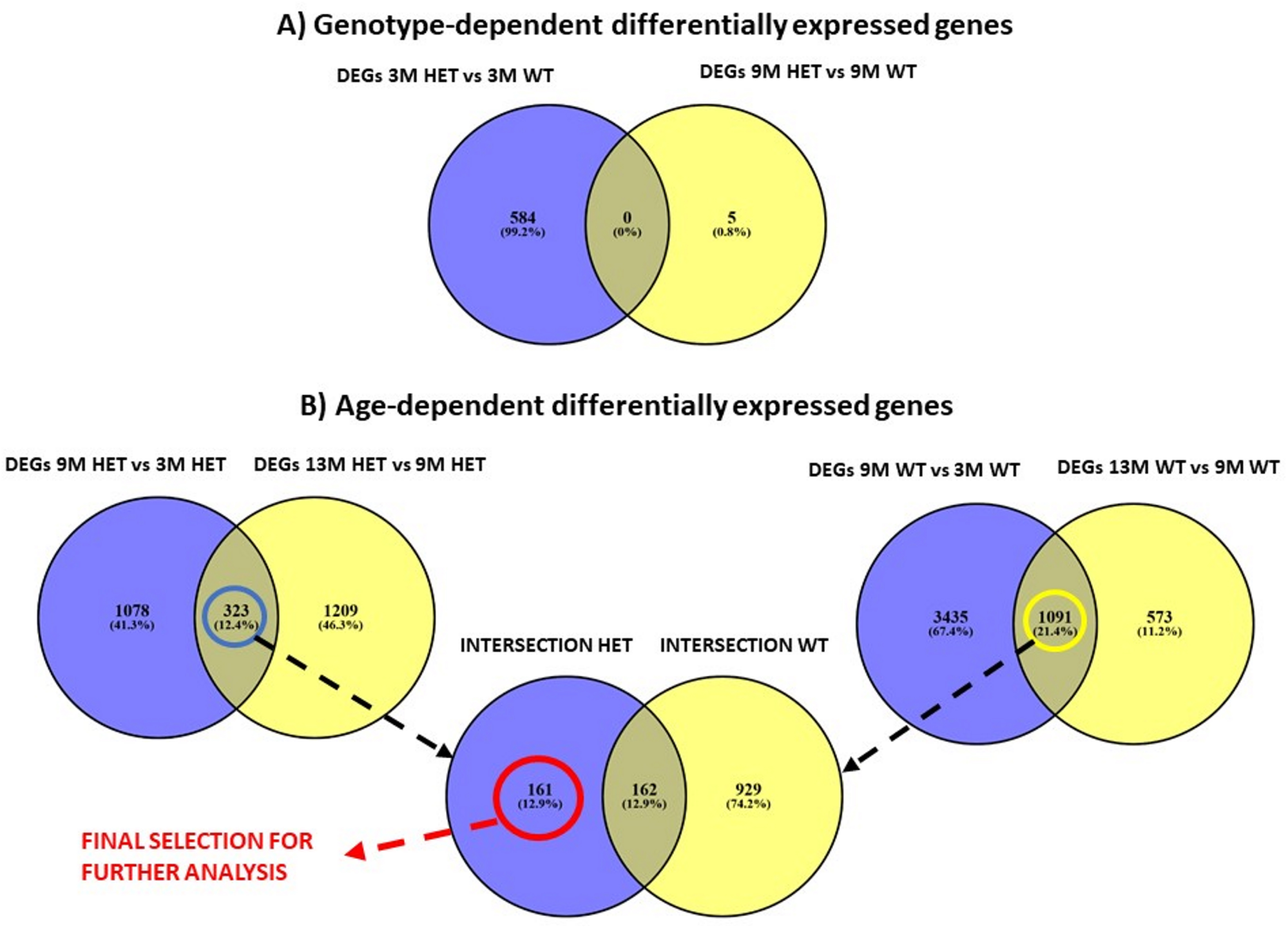
A) Overlap of differentially expressed genes for the age-matched mice of different genotypes (upper panel), B) Differential expression across ages for HET mice (lower panel,left), WT mice (lower panel, right), overlap intersections HET (323 genes) and WT (1091 genes) in previous figures (lower panel, middle).

Then, to uncover age-associated gene expression changes specifically driven by *α*-syn overexpression in our model, we used a stringent selection approach to exclude gene expression changes associated with age in both HET mice and their WT littermates. Our approach is described in the Materials and Methods section, and visualized in the Venn diagrams of Figure 4B.

Since the 9M was the middle age group, we chose this age group, to compare, in each genotype separately, the younger age group (3M) and the older age group (13M) to. Thus for each of the two genotypes, we obtained two sets of DEGs (HET mice: 9M versus 3M: 1401 DEGs, 13M versus 9M: 1532 DEGs; WT mice: 9M versus 3M: 4526 DEGs, 13M versus 9M: 1664 DEGs). For each genotype, these comparisons yielded a significant intersection of DEGs: 323 for HET mice (intersection HET, Fisher’s exact test p-value = 9.27e − 17), and 1091 for WT mice (intersection WT, Fisher’s exact test p-value = 1.78e − 74). These were the genes, for each respective genotype, whose expression changed continuously as the mice aged. To extract the genes whose expression changed continuously over age in HET mice only, we looked at the intersection of the two sets of age-dependent DEGs (intersection HET and intersection WT). The genes whose expression changed continuously over age in HET mice only, a total of 161 DEGs, were selected for further analysis.

#### Genotype-dependent DEGs, their relevance (PMI scores), and cellular sources

Table 2 shows the top 10 DEGs with positive PMI scores in 3M HET mice versus age-matched WT littermates, and the one DEG with a positive PMI score in 9M HET versus age-matched WT littermates, ranked in terms of their PMI text-mining association scores with the MeSH term “Parkinson Disease”.

**Table 2.** Genotype-dependent DEGs (HET vs WT) - DEGs with positive PMI score, ranked in terms of their PMI text-mining association scores with the MeSH term “Parkinson Disease”. Protein names were extracted from Uniprot database (https://www.uniprot.org/) [93]; the brain cell type where the genes show the highest expression was extracted from the Brain RNAseq database (https://www.brainrnaseq.org/) [106].

| age | gene symbol (human) | PMI score | protein name (human) | cell type (human) | gene symbol (mouse) | protein name (mouse) | cell type (mouse) | direction | log2 FC |
| --- | --- | --- | --- | --- | --- | --- | --- | --- | --- |
| 3M (top 10) | TMEM230 | 5.706 | Transmembrane protein 230 | neurons | Tmem230 | Transmembrane protein 230 | oligodendrocytes | up | 0.100 |
|  | RILPL1 | 5.557 | RILP-like protein 1 | astrocytes | Rilpl1 | RILP-like protein 1 | microglia | up | 0.159 |
|  | TMEM175 | 5.403 | Endosomal /lysosomal potassium channel TMEM175 | astrocytes | Tmem175 | Uncharacterized protein | astrocytes | up | 0.076 |
|  | QSER1 | 4.576 | Glutamine and serine-rich protein 1 | astrocytes | Qser1 | Uncharacterized protein | astrocytes | down | -0.120 |
|  | FAM168B | 4.353 | Myelin-associated neurite-outgrowth inhibitor (Mani) (p20) | neurons | Fam168b | Myelin-associated neurite-outgrowth inhibitor | neurons | up | 0.119 |
|  | PSMC4 | 4.016 | 26S proteasome regulatory subunit 6B | microglia | Psmc4 | AAA domain-containing protein | endothelial | up | 0.147 |
|  | IDH3G | 3.765 | Isocitrate dehydrogenase [NAD] subunit, mitochondrial | astrocytes | Idh3g | Isocitrate dehydrogenase [NAD] subunit, mitochondrial | microglia | up | 0.108 |
|  | B4GALNT1 | 3.685 | Beta-1,4-N-acetylgalactosaminyl transferase 1 isoform d | neurons | B4galnt1 | Beta-1,4-N-acetylgalactosaminyl transferase 1 | microglia | up | 0.114 |
|  | COPZ1 | 3.660 | Alternative protein COPZ1 | neurons | Copz1 | Coatomer subunit zeta-1 | microglia | up | 0.109 |
|  | SCAMP5 | 3.660 | Secretory carrier-associated membrane protein | neurons | Scamp5 | Secretory carrier-associated membrane protein | oligodendrocytes | up | 0.079 |
| 9M | OMD | 3.863 | Osteomodulin | endothelial | Omd | Omd protein | pan-cellular | up | 0.113 |
| 13M | no DEGs |  |  |  |  |  |  |  |  |

#### DEGs in 3M HET versus 3M WT

The genotype-dependent DEGs at 3M are expressed across all types of brain cells (Figure 5A). For 30% of DEGs, the main cell type they were expressed in is the same in mouse and human.

**Figure 5.**
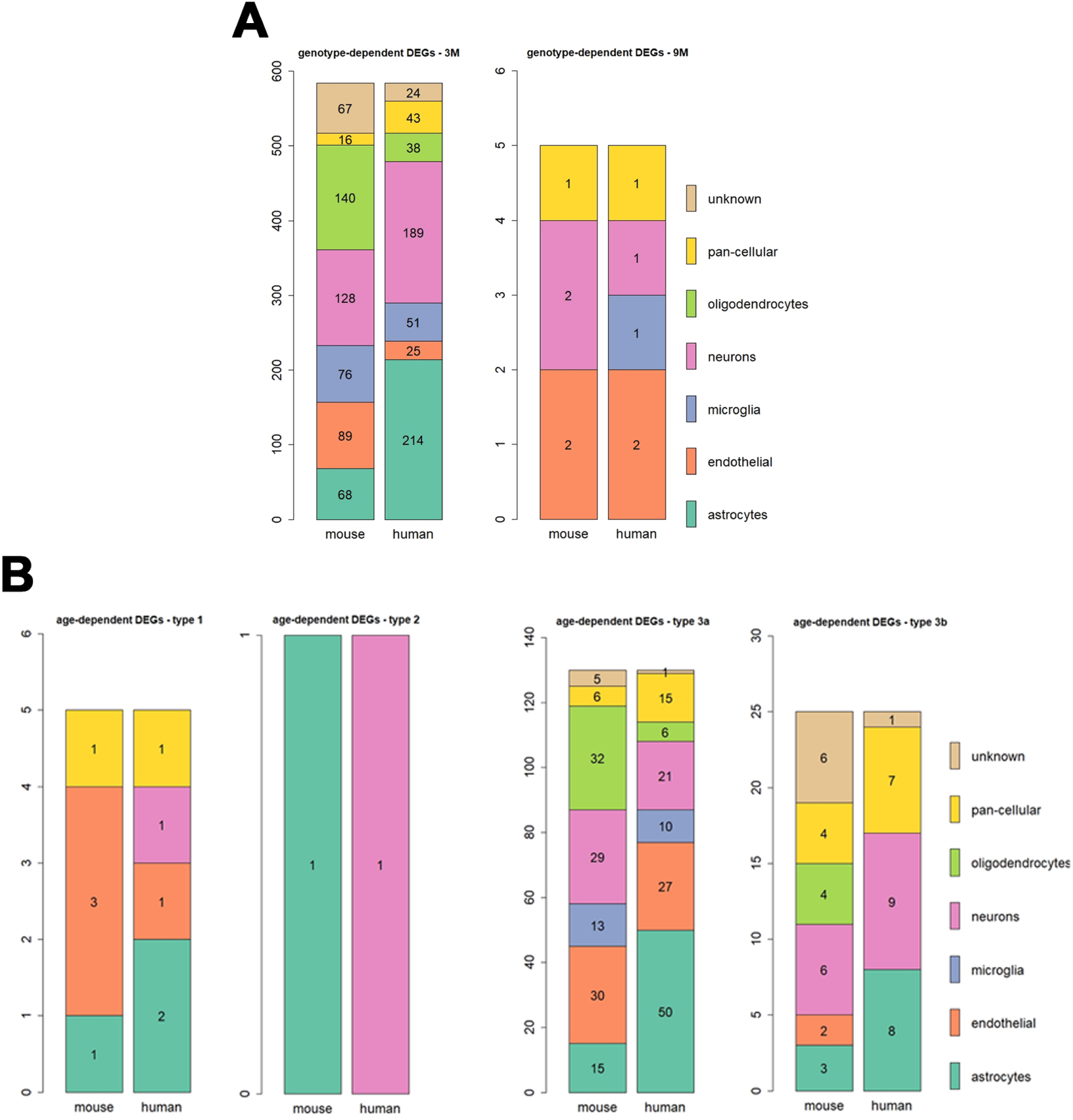
DEGs - brain cell type where the genes show the highest expression (extracted from the Brain RNAseq database (https://www.brainrnaseq.org/) [106]). A) Genotype-dependent DEGs: left panel: 3M (30% same cellular source); right panel: 9M (40% same cellular source). B) Age-dependent DEGs: upper left panel: type 1 - DEGs increasing with age (40% same cellular source); upper right panel: type 2 - DEGs decreasing with age (only 1 DEG, different cellular source between mouse and human); lower left panel: type 3a - DEGs with increased expression between 3M and 9M, and decreased expression between 9M and 13M (28% same cellular source); lower right panel: type 3b - DEGs with decreased expression between 3M and 9M, and increased expression between 9M and 13M *r*(40% same cellular source).

Out of the 584 identified human orthologues, 75 genes had a positive PMI score. In the top 10, 9 were found to be increased in HET versus WT. Four of these genes have previously been associated to *α*-synuclein pathology: *TMEM230* [58], *TMEM175* [41], *B4GALNT1* [96] *and SCAMP5* [100]. *TMEM230* gene product is involved in the trafficking of synaptic vesicles and retromer components [49] and mutations in this gene have been linked to autosomal dominant PD [19]. TMEM175 has been identified as a PD risk locus [70]. It is a K(+) channel regulating lysosomal function [14]. Its deficiency in neuronal cells impairs lysosomal and mitochondrial function, and increases susceptibility to *α*-synuclein toxicity [41]. As an ion channel, it is basically a drugable target. *B4GALNT1* is involved in the synthesis of *GM1* ganglioside, which interacts with *α*-syn to inhibit its fibrilization [96] and promote its clearance [32]. In addition, a PD-like phenotype has been observed in *B4GALNT1* knock-out mice [54]. Thus, the increase of *B4GALNT1* and *TMEM175* in 3M HET mice may indicate an attempt to counteract abnormal *α*-syn. *SCAMP5* has been shown to modulate calcium-dependent exocytosis by interactng with SNAREs [108] and *α*-syn secretion via exomes in vitro [100]. The gene products of two DEGs, measured in blood, have been reported to be associated with PD: *PSMC4* (26S proteasome AAA-ATPase subunit Rpt3, a component of the proteasome complex) [69] and *COPZ1* (COPI coat complex subunit zeta 1, encoding a subunit of the cytoplasmic coatamer protein complex, which is involved in autophagy and intracellular protein trafficking) [79].

#### DEGs in 9M HET versus 9M WT

At 9M, both the number of DEGs and the number of cell types where they are expressed decreased compared to 3M (Figure 5A). For two of the five DEGs, the main cell type where they are expressed is the same in mouse and human. Furthermore, for the human orthologues, microglia appear, whereas these cells do not appear in the mouse chart. The human microglia orthologue (*CRABP2*) is associated with a gene expressed in neurons in mice (*Crabp2*). Of the five DEGs (*OMD,ACTA2, CRABP2, RARB, SLC22A8*) in 9M HET versus 9M WT mice, only *OMD* (Osteomodulin) has a positive PMI score (Table 2, middle part). OMD expression is enhanced in 9M HET mice compared to WT, and its protein level has been shown to be increased in plasma from PD patients, and therefore it could be a potential biomarker [75].

#### DEGs in 13M HET versus 13M WT

No DEG observed.

#### Age-dependent DEGs

Among the 161 age-dependent DEGs being driven by *α*-syn overexpression (Figure 4B, Figure S1, 3 types of expression patterns were observed (see Materials and Methods).

#### Genes with age-dependent increase in expression (Type 1)

These changes may point to key drivers of disease progression, or, conversely, to compensatory and protective mechanisms. Figure 5B shows the brain cell type where these DEGs show the highest expression. When comparing mouse and human, 40% of the DEGs were expressed in the same brain cell type. Of the 5 type 1 DEGs, none has a positive PMI score. The gene product of one of the DEGs though, FAS, has been associated with neurodegeneration in PD, but with some conflicting evidence. One of its ligands which promotes cell death, Fas Associated Factor 1 (FAF1), is the product of a gene that is associated with a form of late-onset PD [35], is increased in PD cortex, and exacerbates the response to PD toxin in vitro [6]. Moreover, the PD-associated mutated Leucine-Rich-Repeat Kinase 2 (LRRK2) was reported to promote neuronal cell death in vitro via a mechanism involving death adaptor FAS-associated protein (FADD), and caspase-8 activation [37]. Absence of FAS partially protected mice against methyl-4-phenyl-1,2,3,6-tetrahydropyridine (MPTP)-induced dopaminergic cell loss in the SN in one study [34], whereas it rendered them more susceptible to it in another [52]. Overall, most of the evidence points to the harming effect of FAS involvement in a PD context, and interfering with the FAS pathway could open up new venues for therapeutic intervention.

#### Genes with age-dependent decrease in expression (Type 2)

These changes may reflect the injury associated with disease progression, and could be potential targets or biomarkers candidates. Only one gene showed decreased expression with age: *UBP1*, a protein coding gene whose product is a transcriptional activator [90]. While for mice this gene is expressed in astrocytes, the human orthologue is expressed in neurons (see Figure 5B). No relationship between *UBP1* and PD has previously been reported in the literature according to the PMI text-mining analysis.

#### Genes switching in directionality with age (Types 3a and 3b)

These DEGs may help to shed some light into more complex adaptive and disease-stage dependent processes involving multiple aspects of disease progression, and will point the way to future studies that help shed light into hitherto unknown facets of PD.

We identified 130 type 3a DEGs that have increased expression between 3M and 9M, then decreased expression between 9M and 13M (Figure S1). 28% of genes are expressed the same main brain cell type in mouse and human. Out of the 130 type 3a DEGs, 18 have a positive PMI score. Table 3 shows the top 10 type 3a DEGs, ranked in terms of their PMI text-mining association scores with the MeSH term “Parkinson Disease”. Among these, *SV2C* modulates dopamine release, and is disrupted in transgenic PD models and in PD, but not in other neurodegenerative diseases [23]. The *ADORA2A* receptor is an GPCR for adenosine, whose disruption is protective in a *α*-syn transgenic model of PD [44], and has been suggested as a potential drug target [15]. Lower expression of *AK4*, a mitochondrial phosphotransferase, has been observed in the late PD [29]. *GPR88*, a Gi/o coupled orphan receptor, has been proposed as a PD target, due to its association with D2 receptor signaling and its involvement in cognitive and motor functions [101]. Finally, *RILPL2* is involved in ciliogenesis linked to downstream *LRRK2* -dependent phosphorylation of Rabs [89]. Mutated forms of *LRRK2* have been reported to impair ciliogenesis as well as its associated neuroprotective Sonic hedgehog signalling [20].

**Table 3.**
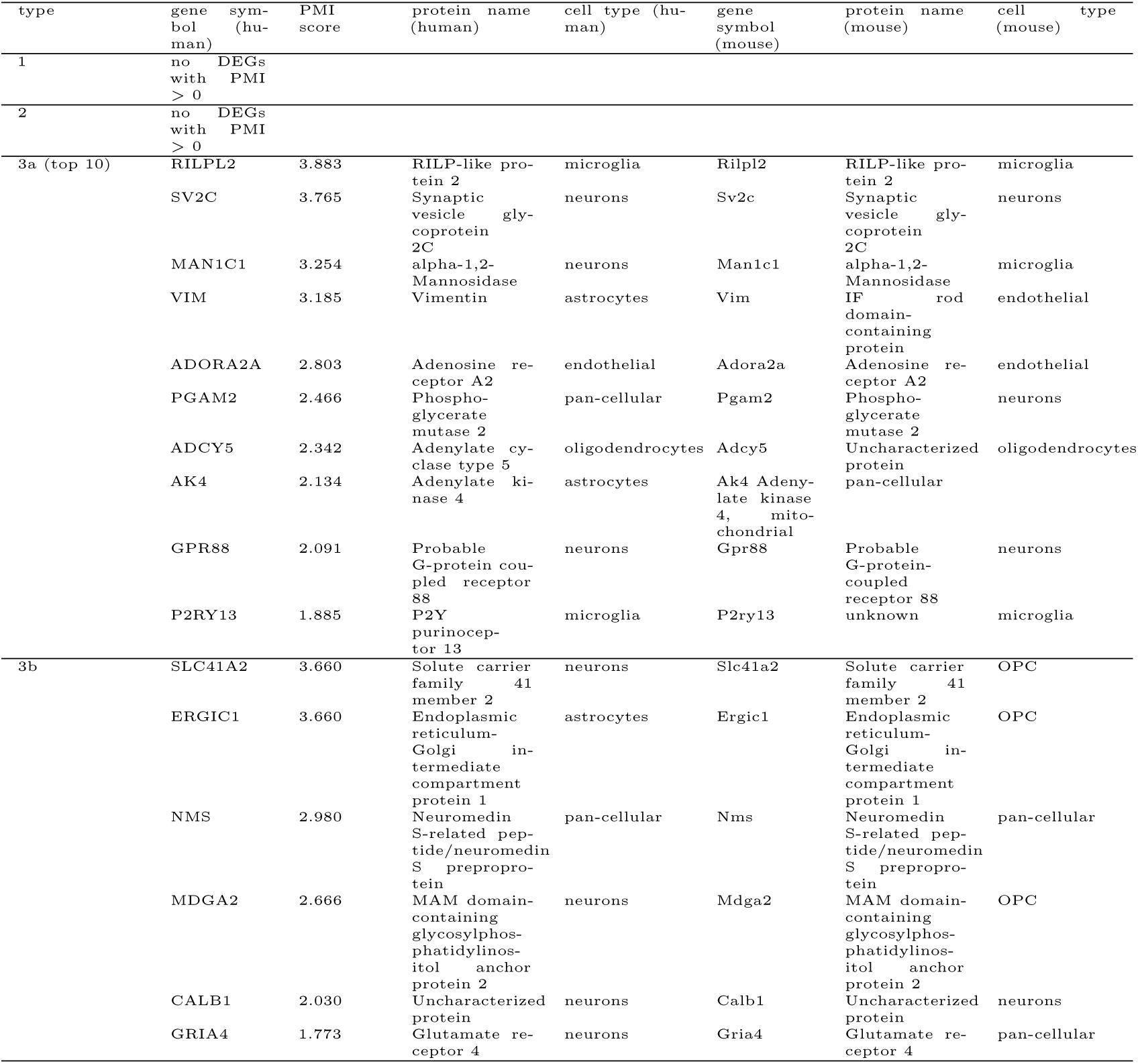
Age-dependent DEGs - DEGs with positive PMI score, ranked in terms of their PMI text-mining association scores with the MeSH term “Parkinson Disease”. Protein names were extracted from Uniprot database (https://www.uniprot.org/) [93]; the brain cell type where the genes show the highest expression was extracted from the Brain RNAseq database (https://www.brainrnaseq.org/) [106]. Type 1: DEGs increasing with age; type 2: DEGs decreasing with age; type 3a: DEGs with increased expression between 3M and 9M, and decreased expression between 9M and 13M; type 3b: DEGs with decreased expression between 3M and 9M, and increased expression between 9M and 13M. OPC: oligodendrocyte progenitor cells.

We identified 25 type 3b genes, that have decreased expression between 3M and 9M, then increased expression between 9M and 13M (Figure S1). 40% of these 3b genes are expressed the same main brain cell type in mouse and human. Like type 3a genes, 3b genes were expressed by various brain cells (Figure 5B). Six out of 25 type 3b DEGs had a positive PMI score (Table 3). Two of these genes have reported association with PD: *MDGA2*, whose product functions as a regulator of axonal growth [42], has a SNP that may be associated with earlier onset in familial PD [53]. *CALB1*, coding for calbinin-1, a protein involved in buffering intracellular calcium, has a SNP associated with sporadic PD [68].

The majority of type 3a and type 3b DEGs were similar in their expression levels in 3M HETs and 13M old HETs, indicating that they transiently increased, or respectively, decreased in 9M old HETs, before returning to the initial expression level. The only exception were 4 type 3a DEGs (*OSR1, SLC22A6, PTN* and *SLC6A20*), whose expression was lower in 13M HETs than in 3 M HETs.

The majority of age-dependent DEGs were switching their expression direction, which was one of the most intriguing observations of our study. We believe this is not an artefact, but a true biological phenomenon. As the mean absolute difference in variance in gene expression between consecutive age groups is very low (Table S2), we can exclude that this observation results from increased variance in the data. Furthermore, we believe that the comparatively lower number of DEGs that increases (type 1) or decreases (type 2) with age in HET mice is also not due to those genes reaching an upward or downward plateau in expression level before the HET mice reach 13M of age. Indeed, the median log2 expression of the genes increasing between 3M and 9M in HET mice was lower than 1 (Figure S2), and, similarly, the median log2 expression of genes decreasing between 3M and 9M in HET mice was only slightly higher than 1 (Figure S2). This indicates that the age-dependent increasing expression of type 1 DEGs does not plateau, and, similarly, age-dependent decreasing expression of type 2 DEGs does not bottom out before HET mice reach 13M. In addition, the 9M old group of HET and WT littermate control mice was derived from the same initial breeding cohorts as their 3M and 13M old counterpart, and the study groups were randomly assigned to the different age groups after weaning and genotyping, thus excluding an inadvertent strain effect due to an error in breeding set-up. Hence, these findings indicate that the age-dependent switch in expression direction of type 3a and type 3b DEGs is a true biological phenomenon that is likely to reflect complex interactions in different responses to the chronic injury in response to *α*-syn overexpression. To the best of our knowledge, such an observation has not been found previously in other PD models.

### Pathway and gene set analysis

Comparative pathway and GO analyses between a disease model and its control reveals molecular and cell biological processes in cells that are altered by disease initiation and progression.

In this study, we were interested in identifying pathways and GO biological processes altered by genotype (SNCA transgene), or changing with age in the context of *α*-syn overexpression in our model.

#### Genotype-dependent pathway/GO changes

##### Pathways/GO biological processes in 3M HET versus 3M WT

Table S3 presents the top 10 out of the 50 identified significantly altered pathways in the 3M HET mice versus the 3M WT mice. Top pathways altered are related to serotonin and glutamate neurotransmissions, which are known to regulate firing activity of SN dopaminergic neurons [66, 107]. Interestingly, a pathway related to G-protein-coupled receptor (GPCR) signaling (“G-protein signaling - Regulation of Cyclic AMP levels by ACM”) was identified. Synucleins (including *α*-syn) are substrates for GPCR kinases [76]. Dopamine D2 receptors, which are present on SN neurons, are also GPCRs [17]. *GPR55* and *GPR37* are GPCR involved in modulating functions of SN neurons, and their absence in mice leads to motor impairments [60, 95]. The prostaglandin (*PGE2*) pathway, an inflammation-related pathway involved in neurodegenerative diseases [56], was identified. Wnt/Beta-catenin and NF-kB pathways were also altered, and are known to modulate dopaminergic cell differentiation and maintenance, as well as to regulate cytokine release from immune cells [61].

Finally, we identified 44 significantly altered GO terms, which (Table S4) included biological processes related to inorganic cation homeostasis, synaptic signaling and memory.

##### Pathways/GO biological processes in 9M HET versus 9M WT

While we found no altered pathway between 9M HETs and their WT littermate controls, we found 66 GO biological processes that differed between them. The top 10 GO terms (Table S4) included developmental processes (e.g. the process “GO:0061448: connective tissue development”). Interestingly, some altered biological processes are associated with regulation of actin filament-based movement, which has been reported to be regulated by *α*-syn [25, 86]. Another altered biological processes related to the ERK1/2 pathway, which is linked to PD pathological processes [10].

##### Pathways/GO biological processes in 13M HET versus 13M WT

Since no DEGs were observed between 13M HETs and their WT littermate controls (see above), no altered pathway or GO were observed either.

#### Age-dependent pathway/GO changes

We used a similar nomenclature (type 1, type 2, type 3a and 3b) for the classification of age-dependent pathway and GO changes as we used for DEGs (see above).

##### Pathways/GO terms with age-dependent upregulation (Type 1)

None identified.

##### Pathways/GO terms with age-dependent downregulation (Type 2)

None identified.

##### Age-dependent pathways/GO terms switching between up/downregulation or vice versa (Types 3a and 3b)

The top 10 out of 50 pathways first upregulated, and then downregulated (type 3a, Table S5) were involved in cell differentiation, migration processes, and GPCR signaling. Interestingly, the pathways identified share signaling components with pathways involved in TGF-beta and PI3K-mediated neuroprotection of dopaminergic neurons [51, 78, 83, 99], and with pathways involved in lipid metabolism and immunomodulation (such as lipoxins [63]). Altered GPCR-related pathways were linked to Nociceptin (*NOP*), C-C chemokine-receptor 1 (*CCR1*), and Sphingosine-1 phosphate-receptor 1 (*S1P1*) signaling. *NOP* is an inhibitory neuropeptide which may promote SN less loss in PD: its genetic deletion protects SN neurons against MPTP in mice, and the inhibition of its receptor protects the same neurons in a rat model of *α*-syn toxicity [3]. *CCR1* is typically microglial and its product participates the recruitment of immune cells, but it is also expressed in the ventral midbrain during development, where it is involved in neuronal differentiation of DA neurons [24]. *S1P1* is uncoupled from an inhibitory G-protein by extracellular *α*-syn, and may therefore play a role in intracellular *α*-syn accumulation [105].

The majority of the top 10 out of the 67 GO terms we detected (Table S6) are involved in GPCR signaling, purinergic signalling, and development.

No pathways, but 117 GO terms first downregulated, then upregulated (type 3b) were identified in HET mice. The top-ranked GO terms are associated with GPCR signaling, calcium homeostasis, and sensory perception (Table S6). The calcium ion homeostasis dysregulation is an important pathological feature of PD [2] and is associated with *α*-syn aggregation and neuronal death [102].

### Network analysis

Molecular networks can provide detailed insights on complex interactions (edges) between biological entities (nodes). In a network analysis of disease-related omics data, the identification of hubs (central nodes) in molecular sub-networks can reveal essential pathological and compensatory responses of the studied disease [1, 40, 71]. For each list of DEGs used for pathway analysis, Table 4 shows the molecular sub-network containing the largest number of input gene products (gene products from DEG list) identified by GeneGO MetaCore*™* ‘s default “Analyze network” procedure.

**Table 4.**
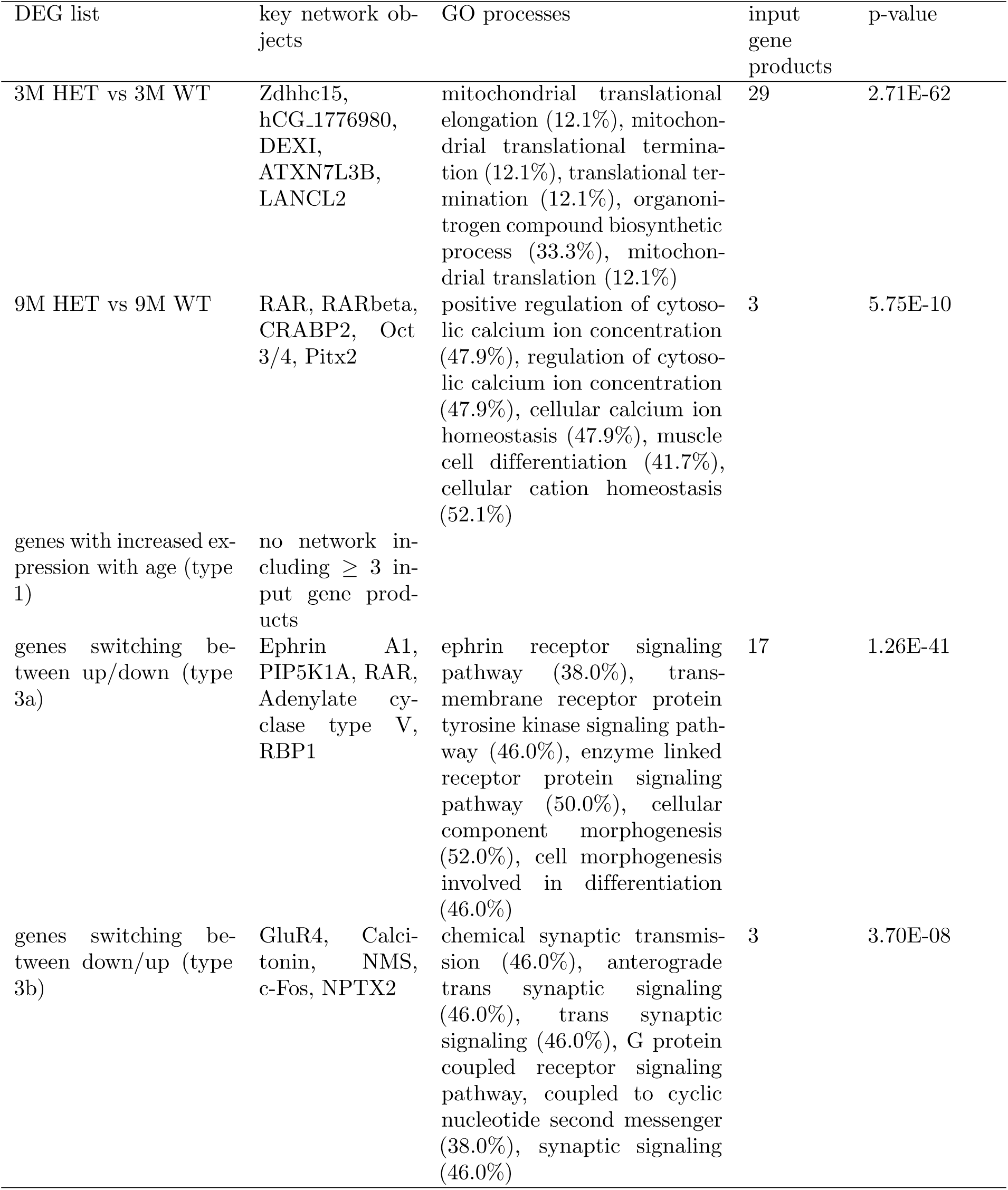
Network with the largest number of input gene products (gene products from DEG list) identified by GeneGO MetaCore*™* ‘s default “Analyze network” procedure. Type 1: 3-9M up, 9-13M up; type 3a: 3-9M up, 9-13M down; type 3b: 3-9M down, 9-13M up.

#### Genotype-dependent networks

In 3M HET mice versus WT littermates, the major sub-network identified (Figure S3) is one involved directly in mitochondrial translation (Table 4). This indicates that mitochondrial dysfunction happens well before striatal neurodegeneration sets in.

In 9 M HET mice versus WT littermates, the altered sub-network (Figure S3) detected is related to calcium ion homeostasis (Table 4). This indicates an active role of calcium dysregulation in the ongoing striatum degeneration.

No network was built for the 13 months old HET vs WT, since no DEGs were identified in that age group.

#### Age-dependent networks

No sub-networks for type 1 DEGs with increased expression with age in HET and containing ≥ 3 input gene products could be identified. No sub-networks for type 2 DEGs with decreased expression with age could be identified.

The type 3a DEGs switching between up/down in HET mice were associated with a sub-network related to cell morphogenesis and differentiation, and Trk signaling (Figure S4, Table 4). These observations are in line with the enriched pathways and GO biological processes shown in Tables S5 and S6.

Finally, for type 3b DEGs switching between down/up in HET mice, the identified sub-network (Figure S4) was involved in synaptic signaling and GPCR signaling, in agreement with the enriched GO terms in Table S6.

### Comparability of the mouse model changes to human PD

When comparing the DEGs obtained in this study to the list of DEGs from a meta-analysis of transcriptomics data from post-mortem brain samples from the SN midbrain region from a previous study [48], we observed a significant overlap with the DEGs between genotypes (HET versus WT) for 3 months old mice (45 genes, Fisher’s exact test p-value = 0.003, see Table S7), but no overlap with the 5 DEGs obtained for 9 months old mice.

Furthermore, also the higher-level analyses of molecular pathways and GO terms point to cellular processes that match with previously proposed mechanisms associated with PD (see previous sections). This, together with the overlap of the genes from a PD meta-analysis, indicates that our model adequately reflects at least a subset of the key molecular features of human PD.

## Discussion

### Mouse models as an essential tool to study early changes in PD

In this study, we provide the first characterization of the novel BAC-Tg3(SNCA*E46K) PD mouse model, a genomic *α*-syn overexpressor carrying the E46K familial PD mutation with a brain regional transgenic expression closely matching that of endogenous *α*-syn. Using a carefully designed bioinformatical analysis pipeline, we investigated the evolution of gene expression profiles, leading up to early PD-like neurodegeneration, and uncovered the underlying molecular events, in this model.

A better understanding of the earliest phases of PD (preclinical, prodromal, and early PD) is essential. Indeed, the consensus is that, in chronic neurodegenerative diseases such as PD, early detection and therapeutic intervention will be instrumental to managing and ultimately, once disease modifying approaches will be available, curing the disease. Getting to this point though is a path paved with obstacles. While deciphering peripheral and non-CNS markers of PD has made some progress [97], understanding what is happening in the brain, in particular the SN, where the degenerating dopaminergic neurons are located, in the earliest phases of the disease, is key to developing early interventions to prevent or limit PD-associated motor dysfunctions.

In studies with that goal, it is important to keep in mind that, for PD or any other brain disease as well as models thereof, the pathological outcomes measured are the result of two different responses to injury: first, the actual degeneration processes, and second, endogenous compensation and/or repair processes. While some measures of molecular pathology clearly indicate one or the other, others can be more challenging to interpret. Also, compensation and repair processes can conceivably, over time, have an effect that is opposite to their original design. One prominent example of this is inflammation which, at least in its initial phases, is designed to clear the organism of infectious agents and damaged tissues, but can end up hurting the organism itself under certain chronic conditions [36].

In PD, the most striking clinical symptoms become obvious only after around 60% of nigral dopaminergic neurons are lost [27], and dopamine concentration in the striatum has declined by around 70% [43]. Thus, it is reasonable to assume that, after PD is initiated and the nigro-striatal circuit starts to be stressed, complex interactions of the two different angles of injury response takes place, possibly over decades, and even before nigro-striatal degeneration occurs. Evidence suggests non-dopaminergic cells (neurons and glia), as well dopaminergic neuron-related processes are contributing to the injury responses of nigral dopaminergic neurons [7, 36, 65].

Several pathological processes such as protein misfolding and abnormal degradation, neuroinflammation, mitochondrial dysfunction, endoplasmatic reticulum stress, and autophagy can all contribute to dopaminergic neuron demise [36, 65]. By contrast, a number of responses of these neurons aimed at limiting dopamine loss or restoring dopaminergic activity have been described [8].

One major obstacle toward a better understanding of the molecular underpinnings of dopaminergic neuron degeneration is the paucity of human brain tissues from the earliest phases of PD, which, for obvious ethical reasons, cannot be obtained routinely. Thus, longitudinal profiling studies on PD animal models, in particular rodents, that develop PD-like phenotypes in an age-dependent fashion, are an important step in shedding light into these complex processes.

We have used one such model, the BAC-Tg3(SNCA^*E*46*K*^) transgenic mouse, to shed light into the molecular events underlying the earliest pathological changes at the gene expression level. This model develops, over age, a phenotype that at least partially resembles early PD. These mice gradually lose TH-positive axonal projections and DAT-positive synapses in their dorsal striatum. We generated longitudinal transcriptional profiles of the ventral midbrain of these mice, a region which contains the dopaminergic neurons that project to the striatum.

### Comparison to previous transcriptomics studies of related rodent models

Only a limited amount of studies have focused so far on transcriptomic profiling in *α*-syn transgenic mouse models of PD. Out of these, only three studies analysed the transcriptomic profile of the midbrain and the dopaminergic neurons therein. Miller et al. [67] analysed microarray data from two *α*-syn transgenic lines, one overexpressing human wildtype *α*-syn, the other one overexpressing a doubly mutated (A30P and A53T) form of *α*-syn, both under the transcriptional control of the rat TH promoter. The 2 lines of mice were profiled both before the onset of neurodegeneration. They found alterations in genes related to dopaminergic neuron function, synaptic function, trophic factors, and protein degradation. Interestingly, similar to our study, they found few inflammation-related genes changes, confirming that, in a PD context, inflammation may be a secondary event, occurring for instance in response to significant neuronal death, or may require a stronger PD-linked pathological challenge than just modest overexpression of *α*-syn. Yacoubian et al. [98] described SN neuron microarray profiling data in an *α*-syn transgenic mice overexpressing human wildtype *α*-syn under the transcriptional control of the pan-neuronal PDGF promoter, before and after the onset of neurodegeneration. Their main findings were gene expression changes of factors involved in DNA transcription, and of some around cell signaling, in their older cohort of mice. Paiva et al. [72] performed RNA-seq on the midbrain of two *α*-syn transgenic lines, one overexpressing human wildtype, the other one A30P *α*-syn, both transgenes being under the transcriptional panneuronal Thy1 promoter. Their most interesting observation was made in the A30P *α*-syn line, where dysregulation of mitochondrial metabolism, endoplasmatic reticulum-related mechanisms, cell adhesion, neuronal development and synaptic activity were observed. The neurodegenerative phenotype of the line overexpressing the A30P mutation is not well known.

One study [13] analysed the transcriptome of the dorsal striatum, the main projections area of SN dopaminergic neurons, of the same *α*-syn overexpressor line as Paiva et al. [72]. Their main observations were changes around synaptic plasticity, neuronal survival and death, signaling, and transcription. Miller et al. [67] also analysed the striatum in their TH-promoter driven wildtype and doubly mutated *α*-syn lines, where they found similar changes. Another line was profiled by these investigators, where transgenic expression of A53T mutated *α*-syn, driven by the prion promoter, leads to a severe age-dependent phenotype in particular in the brainstem and spinal cord. Interestingly, many inflammation-related gene changes could be found, enforcing the notion that strong degeneration may be the driver of an inflammatory response.

There is thus far only one study that applies transcriptomic profiling to a genomic line, similar to the line used here in our study, just that, in that study, the line overexpressed human wildtype *α*-syn, and the investigators profile gene expression in the hippocampus [94]. Most notable changes observed were around synapse function, calcium ion binding, and very generic functions such as “membrane” and “extracellular space”. Interestingly, most of the hippocampal gene expression changes in that line could be prevented by environmental enrichment. An insightful comparative omics profiling study analysed, among other measures, the brain transcriptomic profile of models for different neurodegenerative diseases, including one PD model (Pink-1 knockout mice) [71]. Using RNA extracted from whole brain hemispheres, not from specific disease-susceptible brain regions, the investigators observed that there were few, if any, shared DEGs across different disease models, while still many pathways were common to all the models, suggesting that common pathogenic pathways in neurological disease might not always share the same molecules, or at least not share them in a temporally similar sequence, but still lead to neurodegeneration through shared mechanisms. To the best of our knowledge, no other study has looked so concisely at molecular events leading up to early PD-like neurodegeneration.

### Key observations in the transcriptional profiling of the BAC-Tg3(SNCA^*E*46*K*^) transgenic line

#### Transgenic *α*-syn driven gene expression changes occur early

The vast majority of molecular events linked to transgenic overexpression of human *α*-syn occurred at the youngest age we have looked at (3M), well before striatal degeneration was detectable. The earliest changes were related to mitochondrial translation, followed, at 9M, by calcium homeostasis, intracellular signaling dynamics, and synaptic function. While all these features have been implicated in PD [28], one can assume that, in the early stages they are aimed at fighting off the disease as well as possibly the first signs of neuronal dysfunction. Interestingly, quite few inflammation-related changes were detected. This may be because the microglial function in this and other, similar models, are inhibited as a consequence of the expression of the transgene in those cells [30]. It may also be that stronger disease inducing challenges, like very high *α*-syn overexpression, are needed to start the inflammation process.

#### Number of DEGs and pathways/GO processes decrease with age

Intriguingly, the number of DEGs, and that of their associated pathways and GO biological processes, decreased as the HET mice aged and the striatal degeneration appeared. The mean absolute fold-change between HET and WT mice over all measured genes did also decrease with age, and not due to an increased mean absolute difference in variance between HET and WT (Table S2). This observation is similar to those made in another transgenic *α*-syn model [67], and indicates that the earliest response to the transgenic abnormal *α*-syn challenge is at the gene expression level. Interestingly, Dijkstra et al. [22] reported a lower number of DEGs in SN from patients late Braak stages (3 and up) compared to patients at an early Braak stage (1-2). Similarly, in another study, more altered pathways were observed in incidental Lewy Body Disease compared to PD [80].

While it is possible that these changes are the very first pathological changes, it is tempting to speculate that these are, at least in part, adaptations aimed at compensation that manage to delay onset of striatal degeneration for several months in this model. However, as neurons start to degenerate, their ability to orchestrate a proper gene expression response diminishes, and they start to wither away.

#### The majority of the DEGs were switching their expression directionality, and their associated pathways and GO biological processes switched between up- and downregulation (or vice versa), as the mice aged

This observation, first described in this study, indicates complex interactions between injury versus compensatory responses. A detailed dissection of this phenomenon could lead to novel insights into the PD pathological process, and will be part of future investigations.

#### The cellular sources for DEGs in the different analyses done in this study matched those of their human orthologue counterparts up to 30-40%

Even though we expected more overlap, several points are important to bring up when interpreting this finding. Some gene products, produced by different cells in the two different species, may nonetheless have similar roles in both species. One also has to consider that the database used to determine the cellular source of the DEGs is based on cortical material from mouse and human at baseline, without disease (https://www.brainrnaseq.org/). Under disease conditions, the main cellular source of gene products often shifts. Single-cell profiling studies of midbrain of human PD patients together with similar studies on mouse PD models should shed more light on this potentially confounding issue. Our study stresses for the first time that such considerations should be included when comparing mouse models for neurodegenerative diseases to their human counterparts.

#### Pathway and GO analyses

revealed genotype-dependent alterations in signaling pathways associated with neurotransmission, synaptic plasticity, and lipid metabolism. Lipid metabolism is a feature that has been reported in another *α*-syn-based mouse model [13], and lipids modulate *α*-syn conformation and toxicity [92]. Age-dependent alterations were observed in neuronal and synaptic activity, trans-synaptic signaling and membrane receptor signaling pathways. Importantly, across many of analyses, a dysregulation of GPCR-signaling emerged as a common thread. Since GPCRs are druggable [88], these pathways and their associated DEGs deserve special attention. Dopamine receptors are already a major target of dopamine replacement therapies for PD. Their densities rise in the striatum of PD patients as a compensatory response to loss of dopaminergic neurons in the SN [82]. Our observations indicate that several other GPCRs (see Results) deserve renewed attention.

Overall, our in-depth transcriptome profiling study establishes the BAC-Tg3(SNCA^*E*46*K*^) line as a useful new tool for the investigation of pathological events linked to the development of an early PD-like phenotype, and lays out a detailed set of analysis tools that can be widely applied to uncover similar events in other PD rodent models, or even models for other neurodegenerative diseases.

## Conclusions

In summary, our analyses of a genomic *α*-syn transgenic line have revealed significant molecular changes, in the ventral midbrain, the majority of which precede the appearance of early PD-like striatal neurodegeneration. To the best of our knowledge, this is also the first study on a PD model that lays out a thorough translational assessment by enumerating the human orthologues, and the cellular sources of those genes in both mouse and human, of the disease-associated DEGs and pathways/GO terms observed in different comparisons in our model. It is tempting to speculate that these molecular changes similar to those changes observed in our model happen in midbrain dopaminergic neurons and other cell types in patients well, maybe decades, before any symptoms can be detected. We thus think our model provides a useful new tool for the investigation of the earliest phases of PD.

## Supporting Information

Supplementary figures and tables: see Supplementary material.pdf

## Acknowledgments

The authors also thank Prof. M. Mittelbronn (LCSB/LIH/LNS, Luxembourg), who is funded through a FNR PEARL (P16/BM/11192868) grant, for advice and support. The authors like to thank Klaus Schughart (Helmholtz Center for Infectious Research, Germany) for the mouse breeding.

## Ethics approval

All mouse experiments were performed according to the national guidelines of the animal welfare law in Germany (BGBl. I S. 1206, 1313 and BGBl. I S. 1934). The protocol was reviewed and approved by the ‘Niedersächsisches Landesamt für Verbraucherschutz und Lebensmittelsicherheit, Oldenburg, Germany’ (Permit Number: 33.9-42502-05-11A193).

## Availability of data and materials

The datasets used and analysed during the current study are available from the LCSB (contact: Pierre Garcia) on reasonable request.

