## Supplementary figures and tables for "A new synuclein-transgenic mouse model for early Parkinson’s reveals molecular features of preclinical disease"

March 28, 2020

### Supplementary figures and tables

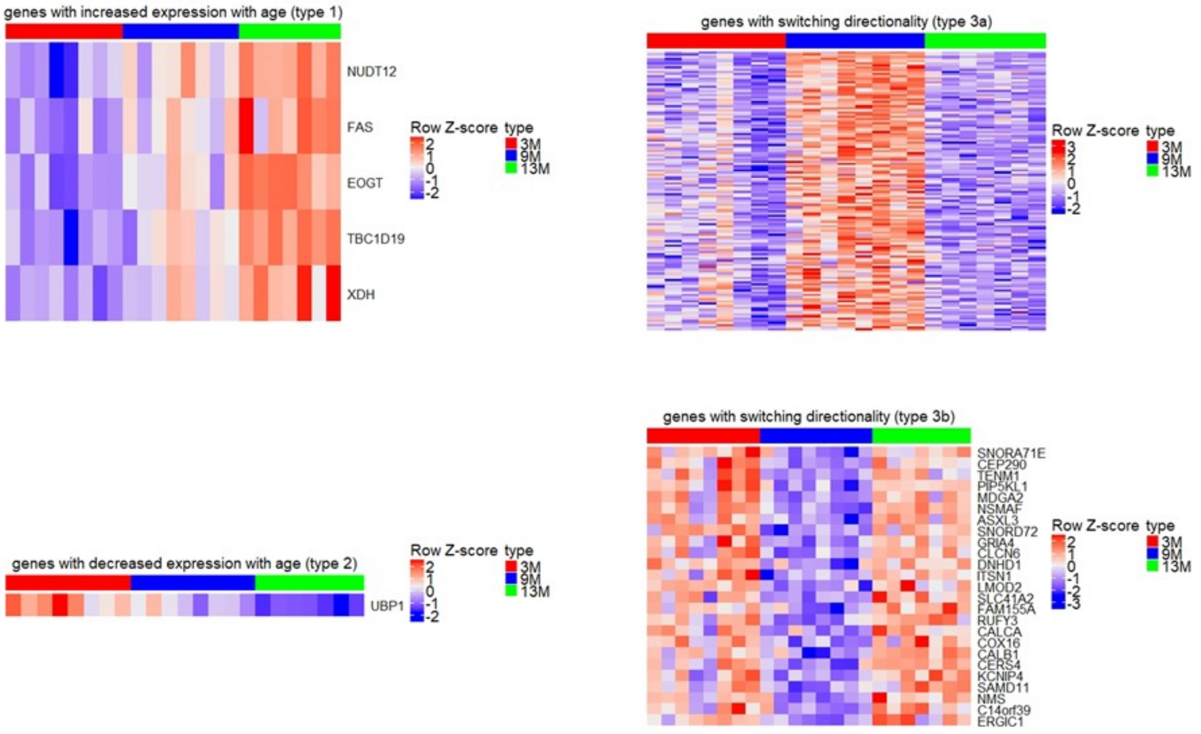


Figure S1: Heat maps (row z-scores) for: genes with increased expression with age (type 1, top left); genes with decreased expression with age (type 2, bottom left); genes with switching directionality up/down (type 3a, top right); genes with switching directionality down/up (type 3b, bottom right).

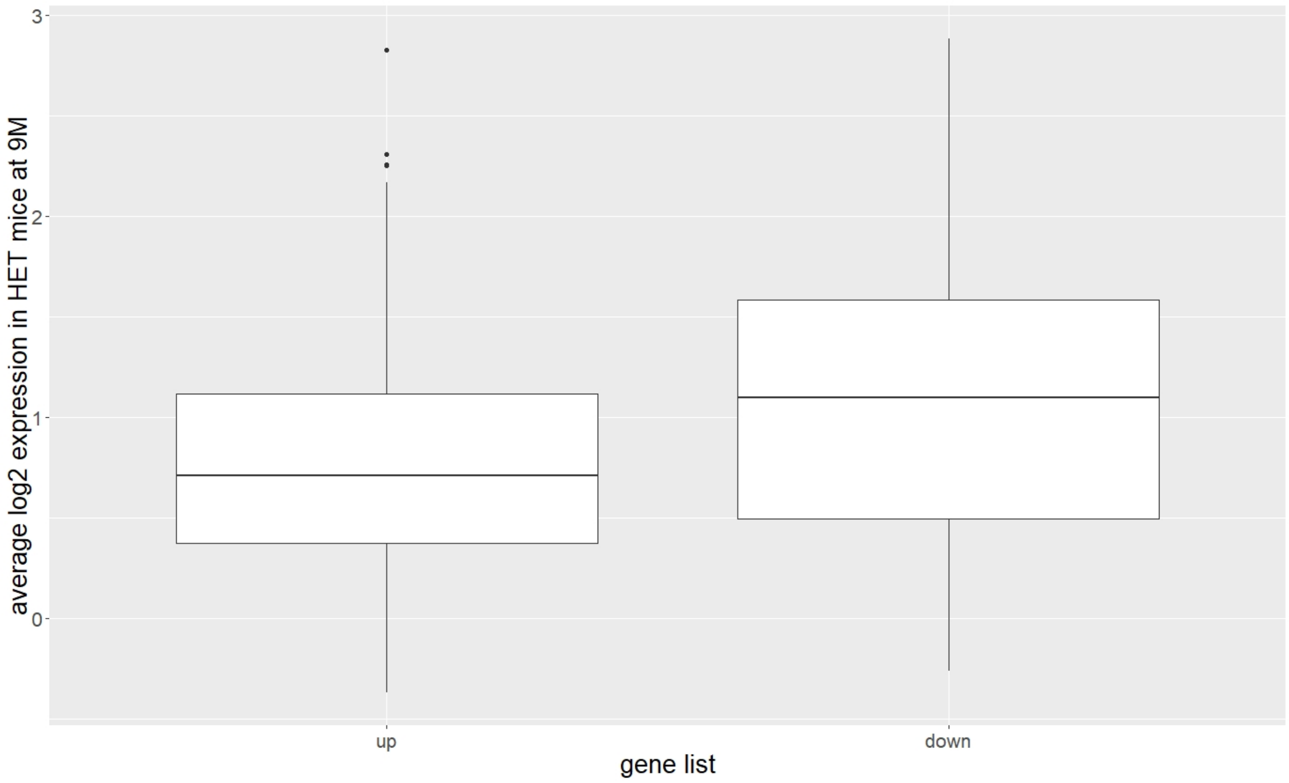


Figure S2: Boxplots of average log2 expression in HET mice at 9M. Left: genes with higher expression between 3M and 9M in HET only, but not differentially expressed between 9M and 13M; right: genes with lower expression between 3M and 9M in HET only, but not differentially expressed between 9M and 13M. The horizontal line in the middle of each boxplot represents the median (50% percentile) expression of the genes within the gene list; the box represents the interquartile range, i.e. the range between the 25% percentile and the 75% percentile; the whiskers represent the lowest and the highest value. Points represent outlying values.

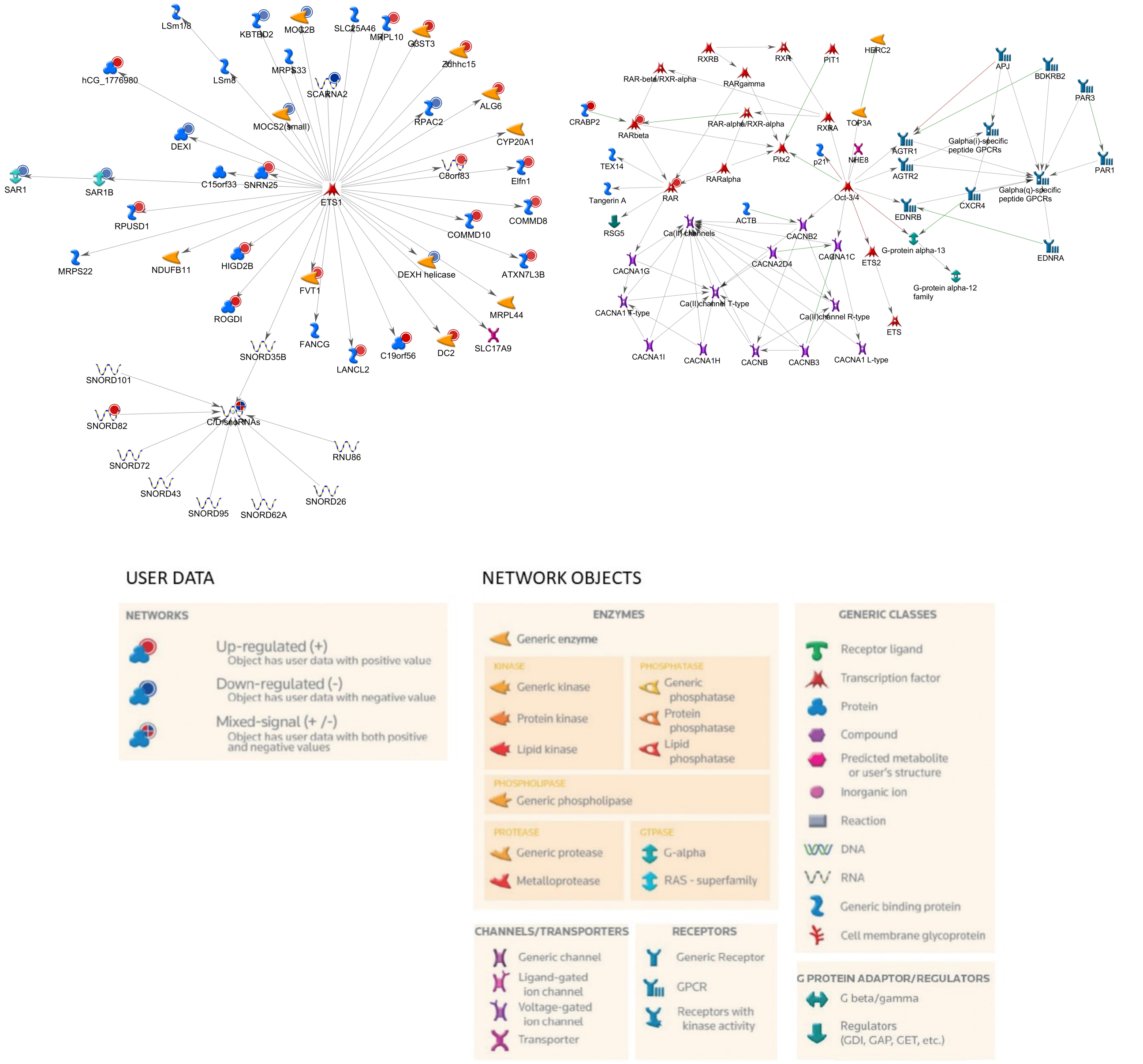


Figure S3: Network with the largest number of input gene products (gene products from DEGs) for 3M HET vs 3M WT, identified by GeneGO MetaCore$^{TM}$'s default "Analyze network" procedure. Gene products of DEGs are indicated with a circle. Left: 3M HET vs 3M WT; right: 9M HET vs 9M WT.

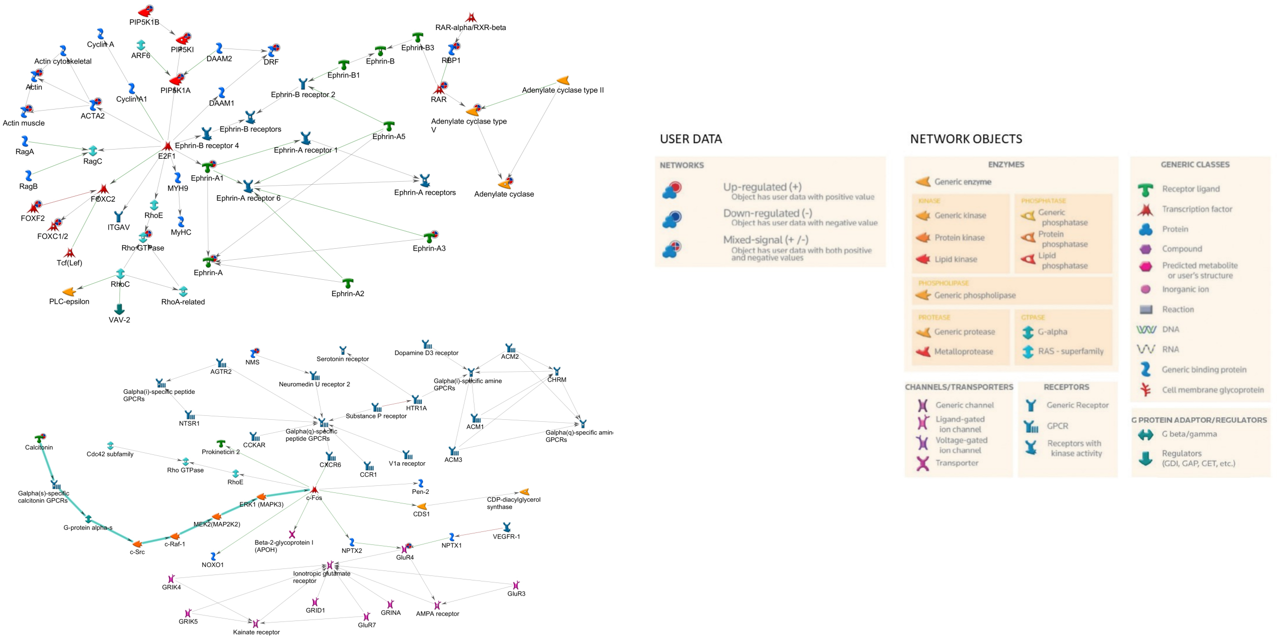


Figure S4: Network with the largest number of input gene products (DEGs) for age-dependent DEGs, identified by GeneGO MetaCore$^{TM}$'s default "Analyze network" procedure. Gene products of DEGs are indicated with a circle. Upper panel: type 3a; lower panel: type 3b. Red-blue: increased expression between 3 months and 9 months, decreased expression between 9 months and 13 months respectively. Cyan line: fragments of canonical pathways.

Table S1: Genes with human orthologues in Ensembl Biomart (`https://www.ensembl.org/biomart`), but classified as mouse specific in NCBI Homologene (`https://www.ncbi.nlm.nih.gov/homologene`). The fourth column indicates in how many databases used by HGNC (`https://www.genenames.org/`) the orthologues show up. The fifth column shows expression in brain in transcripts per million (TPM) according to the Human Proteome Atlas (`https://www.proteinatlas.org/ENSG00000069493-CLEC2D/tissue`).

| mouse | human | DEG list(s) | HGNC | expression in brain (TPM) |
|---|---|---|---|---|
| Tmsb4x | TMSB4Y | 9M HET vs 3M HET, 13M HET vs 9M HET, 9M WT vs 3M WT, 13M WT vs 9M WT | 3 | 3 |
| Rplp1 | RPLP1 | 9M HET vs 3M HET, 9M WT vs 3M WT | 11 | 430 |
| Zfp947 | ZNF34 | 13M HET vs 9M HET, 9M WT vs 3M WT | 1 | 8.4 |
| Josd1 | JOSD1 | 13M HET vs 9M HET | 11 | 39.5 |
| Gm10230 | SYCP3 | 9M WT vs 3M WT | 3 | — |
| Ptma | PTMA | 9M WT vs 3M WT | 7 | 624.3 |
| Gm2030 | SYCP3 | 9M WT vs 3M WT | 3 | — |
| Gm2042 | PRAMEF8 | 9M WT vs 3M WT | 2 | — |
| Gm20738 | SPIN2B | 9M WT vs 3M WT | 1 | 16.1 |
| Vmn1r27 | VN1R5 | 9M WT vs 3M WT | 1 | — |
| Zbed4 | ZBED4 | 9M WT vs 3M WT | 10 | 5.3 |
| Clec2d | CLEC2D | 13M WT vs 9M WT | 6 | 2 |

Table S2: Mean and variance analysis of groups in the dataset. 2nd column: mean absolute logFC (average over all measured genes); 3rd column: mean absolute difference in variance in the data between groups (average over all measured genes).

| **comparison** | **mean absolute logFC** | **mean absolute difference in variance between groups** |
|---|---|---|
| 3M HET vs 3M WT | 0.041 | 0.0042 |
| 9M HET vs 9M WT | 0.029 | 0.0046 |
| 13M HET vs 13M WT | 0.027 | 0.0037 |
| 9M HET vs 3M HET | 0.048 | 0.0042 |
| 13M HET vs 9M HET | 0.053 | 0.0044 |

Table S3: Genotype-dependent pathway alterations (HET vs WT), ranked in terms of their FDR. Fourth column: number of network objects from the data analyzed in this study.

| age | pathway maps | total size | FDR | in DEGs |
|---|---|---|---|---|
| 3M (top 10) | Neurophysiological process - HTR1A receptor signaling in neuronal cells[1] | 43 | 1.816E-03 | 8 |
| | Neurophysiological process - Activity-dependent synaptic AMPA receptor removal | 64 | 1.037E-02 | 8 |
| | Neurophysiological process - Constitutive and regulated NMDA receptor trafficking | 65 | 1.037E-02 | 8 |
| | Development - Epigenetic and transcriptional regulation of oligodendrocyte precursor cell differentiation and myelination | 34 | 1.037E-02 | 6 |
| | Regulation of lipid metabolism - Regulation of lipid metabolism via LXR, NF-Y and SREBP | 38 | 1.528E-02 | 6 |
| | PGE2 pathways in cancer[2] | 55 | 1.528E-02 | 7 |
| | Signal transduction - Angiotensin II/AGTR1 signaling via Notch, Beta-catenin and NF-kB pathways | 76 | 1.683E-02 | 8 |
| | Notch signaling in oligodendrocyte precursor cell differentiation in multiple sclerosis[3] | 27 | 1.987E-02 | 5 |
| | Development - Gastrin in cell growth and proliferation | 62 | 2.306E-02 | 7 |
| | G-protein signaling - Regulation of Cyclic AMP levels by ACM | 45 | 2.379E-02 | 6 |
| 9M | no altered pathways | | | |
| 13M | no altered pathways | | | |

[1] HTR1A: synonym for 5-HT1A

[2] described as cancer pathway in GeneGO MetaCore$^{TM}$, but also an inflammation pathway in neurodegenerative diseases [1]

[3] described as pathway in multiple sclerosis in GeneGO MetaCore$^{TM}$, but also an important pathway in neurodegenerative diseases [2]

Table S4: Genotype-dependent GO biological process (BP) alterations (HET vs WT), ranked in terms of their FDR. Fourth column: number of network objects from the data analyzed in this study.

| age | GO term | total size | FDR | in DEGs |
|---|---|---|---|---|
| 3M (top 10) | GO:0050806: positive regulation of synaptic transmission | 272 | 3.599E-09 | 31 |
| | GO:0055067: monovalent inorganic cation homeostasis | 198 | 1.536E-08 | 25 |
| | GO:0030004: cellular monovalent inorganic cation homeostasis | 159 | 2.710E-08 | 22 |
| | GO:0006885: regulation of pH | 140 | 7.045E-08 | 20 |
| | GO:0007613: memory | 240 | 3.622E-07 | 25 |
| | GO:0030641: regulation of cellular pH | 124 | 1.740E-06 | 17 |
| | GO:0048167: regulation of synaptic plasticity | 327 | 2.183E-06 | 28 |
| | GO:0060251: regulation of glial cell proliferation | 58 | 1.382E-05 | 11 |
| | GO:0045927: positive regulation of growth | 463 | 2.129E-05 | 32 |
| | GO:0098739: import across plasma membrane | 137 | 2.590E-05 | 16 |
| 9M (top 10) | GO:0090131: mesenchyme migration | 12 | 1.272E-06 | 3 |
| | GO:1903116: positive regulation of actin filament-based movement | 16 | 1.527E-06 | 3 |
| | GO:1904238: pericyte cell differentiation | 17 | 1.527E-06 | 3 |
| | GO:0061448: connective tissue development | 343 | 3.948E-06 | 5 |
| | GO:0002064: epithelial cell development | 350 | 4.093E-06 | 5 |
| | GO:0072132: mesenchyme morphogenesis | 72 | 5.986E-05 | 3 |
| | GO:1903115: regulation of actin filament-based movement | 79 | 7.446E-05 | 3 |
| | GO:0070374: positive regulation of ERK1 and ERK2 cascade | 369 | 1.397E-04 | 4 |
| | GO:0071300: cellular response to retinoic acid | 134 | 2.680E-04 | 3 |
| | GO:0070372: regulation of ERK1 and ERK2 cascade | 481 | 3.312E-04 | 4 |
| 13M | no altered GO BP | | | |

Table S5: Age-dependent pathway alterations, ranked in terms of their FDR. Fourth column: number of network objects from the data analyzed in this study. Type 1: DEGs increasing with age; type 2: DEGs decreasing with age; type 3a: DEGs with increased expression between 3M and 9M, and decreased expression between 9M and 13M; type 3b: DEGs with decreased expression between 3M and 9M, and increased expression between 9M and 13M.

| type | pathway maps | total size | FDR | in DEGs |
|---|---|---|---|---|
| type 1 | no pathway alterations | | | |
| type 2 | no pathway alterations | | | |
| type 3a (top 10) | Development - Astrocyte differentiation from adult stem cells | 40 | 1.987E-04 | 6 |
| | Chemotaxis - Inhibitory action of lipoxins on IL-8- and Leukotriene B4-induced neutrophil migration | 53 | 7.330E-04 | 6 |
| | Nociception - Nociceptin receptor signaling | 76 | 2.987E-03 | 6 |
| | Development - TGF-beta-dependent induction of EMT via RhoA, PI3K and ILK | 46 | 2.987E-03 | 5 |
| | Development - S1P2 and S1P3 receptors in cell proliferation and differentiation | 26 | 3.015E-03 | 4 |
| | Chemotaxis - CCR1 signaling | 53 | 3.241E-03 | 5 |
| | Impaired inhibitory action of lipoxins on neutrophil migration in CF | 56 | 3.912E-03 | 5 |
| | Development - S1P1 receptor signaling via beta-arrestin | 34 | 5.460E-03 | 4 |
| | Stem cells - Response to hypoxia in glioblastoma stem cells | 40 | 9.357E-03 | 4 |
| | Stem cells - H3K27 demethylases in differentiation of stem cells | 40 | 9.357E-03 | 4 |
| type 3b | no pathway alterations | | | |

Table S6: Age-dependent GO biological process (BP) alterations, ranked in terms of their FDR. Fourth column: number of network objects from the data analyzed in this study. Type 1: DEGs increasing with age; type 2: DEGs decreasing with age; type 3a: DEGs with increased expression between 3M and 9M, and decreased expression between 9M and 13M; type 3b: DEGs with decreased expression between 3M and 9M, and increased expression between 9M and 13M.

| type | GO term | total size | FDR | in DEGs |
|---|---|---|---|---|
| type 1 | no altered GO BP | | | |
| type 2 | no altered GO BP | | | |
| type3a (top 10) | GO:0035588: G protein-coupled purinergic receptor signaling pathway | 29 | 7.908E-11 | 9 |
| | GO:0035587: purinergic receptor signaling pathway | 38 | 6.608E-10 | 9 |
| | GO:0001973: adenosine receptor signaling pathway | 15 | 6.608E-10 | 7 |
| | GO:0061448: connective tissue development | 343 | 3.339E-09 | 18 |
| | GO:0072132: mesenchyme morphogenesis | 72 | 6.974E-09 | 10 |
| | GO:0050768: negative regulation of neurogenesis | 498 | 1.978E-08 | 20 |
| | GO:0031279: regulation of cyclase activity | 86 | 3.030E-08 | 10 |
| | GO:0006171: cAMP biosynthetic process | 17 | 7.385E-08 | 6 |
| | GO:0045665: negative regulation of neuron differentiation | 403 | 1.688E-07 | 17 |
| | GO:0007188: adenylate cyclase-modulating G protein-coupled receptor signaling pathway | 306 | 1.914E-07 | 15 |
| type 3b (top 10) | GO:1990408: calcitonin gene-related peptide receptor signaling pathway | 9 | 2.861E-05 | 3 |
| | GO:0032730: positive regulation of interleukin-1 alpha production | 10 | 3.063E-05 | 3 |
| | GO:0097647: amylin receptor signaling pathway | 13 | 4.629E-05 | 3 |
| | GO:0097646: calcitonin family receptor signaling pathway | 14 | 4.629E-05 | 3 |
| | GO:0007218: neuropeptide signaling pathway | 146 | 4.629E-05 | 5 |
| | GO:0032650: regulation of interleukin-1 alpha production | 15 | 4.629E-05 | 3 |
| | GO:0045651: positive regulation of macrophage differentiation | 23 | 1.493E-04 | 3 |
| | GO:0002031: G protein-coupled receptor internalization | 25 | 1.660E-04 | 3 |
| | GO:0050961: detection of temperature stimulus involved in sensory perception | 31 | 2.826E-04 | 3 |
| | GO:0050965: detection of temperature stimulus involved in sensory perception of pain | 31 | 2.826E-04 | 3 |

Table S7: Overlap DEGs 3M HET vs 3M WT with DEGs from a recent meta-analysis on PD in human [3]. Protein names were extracted from Uniprot database (https://www.uniprot.org/) [4]; the brain cell type where the genes show the highest expression was extracted from the Brain RNAseq database (https://www.brainrnaseq.org/) [5].

| gene symbol (human) | protein name (human) | cell type (human) | gene symbol (mouse) | protein name (mouse) | cell type (mouse) |
|---|---|---|---|---|---|
| OLFM1 | Noelin | neurons | Olfm1 | Noelin | neurons |
| KIF3C | Kinesin-like protein KIF3C | neurons | Kif3c | Kinesin-like protein | neurons |
| NLK | Mitogen-activated protein kinase | neurons | Nlk | Serine/threonine-protein kinase NLK | endothelial |
| SLC25A16 | Graves disease carrier protein | neurons | Slc25a16 | Graves disease carrier protein homolog | astrocytes |
| SCAMP1 | Secretory carrier-associated membrane protein 1 | neurons | Scamp1 | Secretory carrier-associated membrane protein | neurons |
| GNG3 | Guanine nucleotide-binding protein G(I)/G(S)/G(O) subunit gamma-3 | neurons | Gng3 | Guanine nucleotide-binding protein G(I)/G(S)/G(O) subunit gamma-3 | neurons |
| CXXC1 | CXXC-type zinc finger protein 1 | astrocytes | Cxxc1 | Uncharacterized protein | endothelial |
| SNX10 | Sorting nexin-10 | neurons | Snx10 | PX domain-containing protein | neurons |
| TUBA4A | Tubulin alpha chain | neurons | Tuba4a | Tubulin alpha-4A chain | microglia |
| STAU2 | Double-stranded RNA-binding protein Staufen homolog 2 | neurons | Stau2 | Double-stranded RNA-binding protein Staufen homolog 2 | neurons |
| LMO3 | LIM domain only protein 3 | astrocytes | Lmo3 | LIM domain only protein 3 | unknown |
| TUSC2 | Tumor suppressor candidate 2 | neurons | Tusc2 | Tumor suppressor candidate 2 | OPC |
| ATP6V1C1 | V-type proton ATPase subunit C | neurons | Atp6v1c1 | V-type proton ATPase subunit C | microglia |
| ARHGEF9 | Rho guanine nucleotide exchange factor 9 | neurons | Arhgef9 | Rho guanine nucleotide exchange factor 9 | neurons |
| SLC25A22 | Mitochondrial glutamate carrier 1 | astrocytes | Slc25a22 | Mitochondrial glutamate carrier 1 | neurons |
| PCDH8 | Protocadherin-8 | neurons | Pcdh8 | Protocadherin-8 | neurons |
| LIN7B | Protein lin-7 homolog B | endothelial | Lin7b | Protein lin-7 homolog B | neurons |
| SNAPC5 | snRNA-activating protein complex subunit 5 | astrocytes | Snapc5 | snRNA-activating protein complex subunit 5 | endothelial |
| SEH1L | Nucleoporin SEH1 | neurons | Seh1l | Nucleoporin SEH1 | neurons |
| ATXN7L3B | Ataxin-7-like protein 3B | neurons | Atxn7l3b | Ataxin-7-like protein 3B | pan-cellular |
| FBXO21 | F-box only protein 21 | neurons | Fbxo21 | Uncharacterized protein | neurons |
| CDH19 | Cadherin-19 | oligodendrocytes | Cdh19 | Uncharacterized protein | oligodendrocytes |
| PKNOX2 | Homeobox protein PKNOX2 | neurons | Pknox2 | Meis_PKNOX_N domain-containing protein | neurons |
| GOT1 | Aspartate aminotransferase, cytoplasmic | pan-cellular | Got1 | Aspartate aminotransferase, cytoplasmic | pan-cellular |
| PID1 | PTB-containing, cubilin and LRP1-interacting protein | neurons | Pid1 | Uncharacterized protein | OPC |
| TOMM20 | Mitochondrial import receptor subunit TOM20 homolog | neurons | Tomm20 | Mitochondrial import receptor subunit TOM20 homolog | endothelial |
| CDC27 | Cell division cycle protein 27 | neurons | Cdc27 | Uncharacterized protein | endothelial |
| ATP6V1E1 | V-type proton ATPase subunit E 1 | neurons | Atp6v1e1 | V-type proton ATPase subunit E 1 | unknown |
| PLD3 | Phospholipase D3 | astrocytes | Pld3 | Phospholipase D3 | microglia |
| ATP1A1 | Sodium/potassium-transporting ATPase subunit alpha-1 | neurons | Atp1a1 | Sodium/potassium-transporting ATPase subunit alpha | endothelial |
| B4GALT6 | Beta-1,4-galactosyltransferase 6 | neurons | B4galt6 | Uncharacterized protein | neurons |
| SCAMP5 | Secretory carrier-associated membrane protein | neurons | Scamp5 | Secretory carrier-associated membrane protein | oligodendrocytes |
| SMIM14 | Small integral membrane protein 14 | neurons | Smim14 | Small integral membrane protein 14 | oligodendrocytes |
| RNF10 | RING finger protein 10 | astrocytes | Rnf10 | RING finger protein 10 | OPC |
| MAPK1 | Mitogen-activated protein kinase | neurons | Mapk1 | Mapk1 protein | neurons |
| SLC35F1 | Solute carrier family 35 member F1 | neurons | Slc35f1 | Solute carrier family 35 member F1 | OPC |
| KANSL2 | KAT8 regulatory NSL complex subunit 2 | neurons | Kansl2 | KAT8 regulatory NSL complex subunit 2 | OPC |
| PELI1 | E3 ubiquitin-protein ligase pellino homolog 1 | microglia | Peli1 | E3 ubiquitin-protein ligase pellino homolog | oligodendrocytes |
| CHD7 | Chromodomain-helicase-DNA-binding protein 7 | astrocytes | Chd7 | Chromodomain-helicase-DNA-binding protein 7 | OPC |
| MAN1C1 | alpha-1,2-Mannosidase | neurons | Man1c1 | alpha-1,2-Mannosidase | microglia |
| COPS8 | COP9 signalosome complex subunit 8 | neurons | Cops8 | COP9 signalosome complex subunit 8 | OPC |
| KDSR | 3-ketodihydrosphingosine reductase | oligodendrocytes | Kdsr | Uncharacterized protein | oligodendrocytes |
| GMEB2 | Alternative protein GMEB2 | astrocytes | Gmeb2 | Glucocorticoid modulatory element-binding protein 2 | astrocytes |
| RPL38 | 60S ribosomal protein L38 | oligodendrocytes | Rpl38 | 60S ribosomal protein L38 | endothelial |
| SNRNP25 | U11/U12 small nuclear ribonucleoprotein 25 kDa protein | neurons | Snrnp25 | U11/U12 small nuclear ribonucleoprotein 25 kDa protein | oligodendrocytes |
